# Different tertiary interactions create the same important 3-D features in a divergent flavivirus xrRNA

**DOI:** 10.1101/2020.06.25.172429

**Authors:** Rachel A. Jones, Anna-Lena Steckelberg, Matthew J. Szucs, Benjamin M. Akiyama, Quentin Vicens, Jeffrey S. Kieft

**Author notes:** To whom correspondence should be addressed: Jeffrey S. Kieft, Department of Biochemistry and Molecular Genetics, University of Colorado Denver School of Medicine, Mail Stop 8101, Aurora, CO 80045.

## Abstract

During infection by a flavivirus (FV), cells accumulate noncoding subgenomic flavivirus RNAs (sfRNAs) that interfere with several antiviral pathways. These sfRNAs are formed by structured RNA elements in the 3′ untranslated region (UTR) of the viral genomic RNA, which block the progression of host cell exoribonucleases that have targeted the viral RNA for destruction. Previous work on these exoribonuclease-resistant RNAs (xrRNAs) from mosquito-borne FVs revealed a specific 3-dimensional fold with a unique topology in which a ring-like structure protectively encircles the 5′ end of the xrRNA. Conserved nucleotides make specific tertiary interactions that support this fold. Examination of more divergent FVs reveals differences in their 3′ UTR sequences, raising the question of whether they contain xrRNAs and if so, how they fold. To answer this, we demonstrated the presence of an authentic xrRNA in the 3′ UTR of the Tamana Bat Virus (TABV) and solved its structure by x-ray crystallography. The structure reveals conserved features from previously characterized xrRNAs, but in the TABV version these features are created through a novel set of tertiary interactions not previously seen in xrRNAs. This includes two important A-C interactions, four distinct backbone kinks, several ordered Mg^2+^ ions, and a C^+^-G-C base triple. The discovery that the same overall architecture can be achieved by very different sequences and interactions in distantly related flaviviruses provides insight into the diversity of this type of RNA and will inform searches for undiscovered xrRNAs in viruses and beyond.

## INTRODUCTION

Members of the *Flaviviridae* family of viruses are major threats to human health. Approximately 2/3^rd^ of the world’s population are at risk for infection by one of the 35 different disease-causing flaviviruses, which include Dengue, West Nile, Zika, Japanese encephalitis, and yellow fever viruses (Fields et al. 2013). Flavivirus genomes are single-stranded positive-sense RNAs ∼9.6 kilobases long that contain a single open reading frame flanked by 5′ and 3′ untranslated regions (UTRs). These UTRs contain a number of structured RNA elements that play critical roles during infection, to include coordinating and regulating replication and translation (Khromych 2001/(Selisko et al. 2014; Brinton and Basu 2015; Ng et al. 2017). During infection, the genomic RNA is replicated and accumulates in the cell, but several *Flaviviridae* clades also produce high levels of another 300-500 nucleotide-long RNA species called ‘subgenomic flaviviral RNAs’ (sfRNAs), which are non-coding RNAs derived from the 3’UTR of the viral genomic RNA(Naeve and Trent 1978; Takeda et al. 1978; Wengler et al. 1978; Clarke et al. 2015; Slonchak and Khromykh 2018).

sfRNAs have been associated with pathogenic outcomes and cytopathicity during infection (Funk et al. 2010; Chang et al. 2013; Walker et al. 2013; Donald et al. 2016; Filomatori et al. 2017; Junglen et al. 2017). Although their full set of functions remains an active area of investigation (Slonchak and Khromykh 2018) sfRNAs have been shown to bind a number of host proteins (Bidet et al. 2014; Schnettler et al. 2014; Manokaran et al. 2015; Moon et al. 2015; Goertz et al. 2019; Michalski et al. 2019) suggesting they affect several processes during infection including inhibiting the interferon response (Schuessler et al. 2012) and suppressing RNAi pathways in insects (Schnettler et al. 2012; Schnettler et al. 2014; Moon et al. 2015). Furthermore, there is evidence that sfRNAs are involved in switching between insect vectors and mammalian hosts (Filomatori et al. 2017; de Borba et al. 2019), enhancing transmission by mosquitoes (Pompon et al. 2017; Yeh and Pompon 2018; Goertz et al. 2019; Slonchak et al. 2020), and altering the cell’s mRNA decay program (Moon et al. 2012; Michalski et al. 2019). The importance of sfRNAs makes it critical to understand how they are produced.

sfRNAs form through adventitious use of the cell’s normal cytoplasmic messenger RNA (mRNA) decay pathway. Specifically, a subset of the viral genomic RNA is targeted for degradation by host cell exoribonuclease Xrn1 (Jones et al. 2012; Nagarajan et al. 2013), which loads on the 5′ end of either decapped or endonucleolytically-cleaved 5’ monophosphorylated viral RNA and processively degrades it in a 5′ to 3′ direction. Upon reaching specifically-structured elements in the 3′UTR, Xrn1 halts and the RNA that is protected from degradation becomes an sfRNA (Pijlman et al. 2008; Funk et al. 2010; Silva et al. 2010; Moon et al. 2012; Chapman et al. 2014a; Chapman et al. 2014b; Akiyama et al. 2016; MacFadden et al. 2018)(**Fig. 1A**). The elements in the viral 3′ UTR that block the enzyme are called exoribonuclease-resistant RNAs (xrRNAs), because they block progression of the nuclease using a discrete RNA structure without needing accessory proteins (Chapman et al. 2014b).

**Figure 1.**
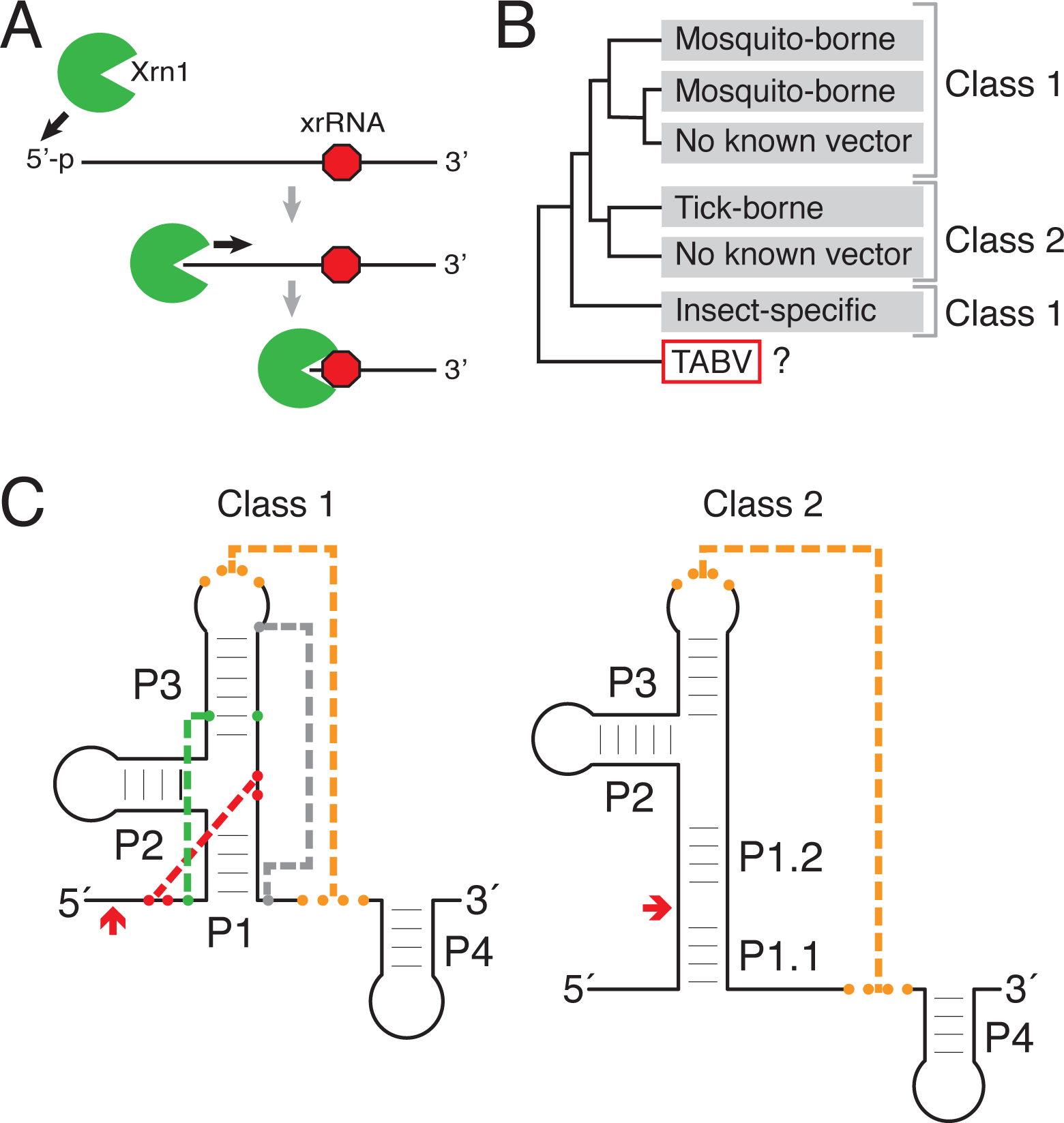
Function, classification, and distribution of known xrRNAs in *Flaviviridae*. (A) xrRNA elements form structures that block 5 τo 3′ degradation by Xrn1 (green) at a defined location (red octagon) without requiring bound accessory proteins. Here, only a single halt point is shown; some viral RNAs contain multiple xrRNAs (MacFadden et al. 2018). (B) Schematic of the phylogenetic relationship of genera within the *Flaviviridae* (Shi et al. 2018) and associated xrRNA classes. (C) xrRNA classes represented as cartoons with secondary and tertiary structure elements labeled. Orange: long-range pseudoknot; red: internal pseudoknot; green: base triple; grey: the long-range interaction that closes the ring-like structure. Red arrows denote the approximate halt site of Xrn1.

xrRNAs have now been identified in several genera of the *Flaviviridae* family, but most three-dimensional structural knowledge is derived from mosquito-borne flavivirus (MBFV) examples. Structures of MBFVs xrRNAs solved by x-ray crystallography revealed a fold with a unique topology in which a ring-like structure wraps protectively around the 5′ end of the xrRNA (Chapman et al. 2014a; Kieft et al. 2015; Akiyama et al. 2016). Mechanistic studies suggest a model in which this fold acts as a molecular brace that physically blocks the progression of Xrn1 past a defined point, in fact these xrRNAs can block a variety of exoribonucleases from diverse sources (MacFadden et al. 2018). This ring-like structure and mechanism was also observed in an xrRNA from a plant-infecting dianthovirus and is now known to be widespread in several plant viral families, although the structural strategy used to form the ring-like structure differs from the MBFVs’ (Iwakawa et al. 2008; Steckelberg et al. 2018a; Steckelberg and Vicens et al. 2018b; Steckelberg and Vicens et al. 2020). Thus, different structural strategies can create exoribonuclease-resistant elements, motivating efforts to understand the full diversity of xrRNA structure.

Within the *Flaviviridae* family, two classes of xrRNAs have been identified (**Fig. 1B&C**). The flavivirus class 1 xrRNAs thus far exist in the MBFVs, a few no known vector viruses (NKV) and potentially the insect-specific flaviviruses (ISVs) (MacFadden et al. 2018). Detailed three-dimensional tertiary structure information and functional studies coupled with mutagenesis show that the unique ring-like fold in the class 1 requires a specific set of long-range tertiary contacts (Chapman et al. 2014a; Kieft et al. 2015; Akiyama et al. 2016); and a distinct primary, secondary, and tertiary structure conservation pattern defines this xRNA class (**Fig. 1C, Sup. Fig. S1A**). The flavivirus class 2 xrRNAs are associated with the tick-borne flavivirus (TBFV) and some NKV (MacFadden et al. 2018). Although tertiary structure-based analysis of the class 2 is not yet possible due to a lack of three-dimensional structural information, several sequence and secondary structure patterns present in all class 1 are not found in these class 2. Importantly, assignment of xrRNAs from different clades to class 1 or class 2 should not be considered final, as the discovery of new viruses and new xrRNAs may reveal exceptions.

The discovery of new and diverse *Flaviviridae* has continued to provide new sequence information, allowing further analysis of xrRNA structure diversity. These discoveries have revealed potential xrRNAs that are not readily placed into either the class 1 or class 2. An example is found in the Tamana bat virus (TABV), a virus with no known arthropod vector and classified as a tentative species in the *Flavivirus* genus because it is considered highly divergent (de Lamballerie et al. 2002; Fauquet et al. 2005; Maruyama et al. 2014). Examination of the 3′ UTR of TABV revealed structures that could potentially be xrRNAs (Ochsenreiter et al. 2019), but they lack the important conserved sequences required for forming the Xrn1-resistance fold in the MBFV class 1 xrRNAs and definitive assignment to either class was ambiguous.

To better understand the diversity of xrRNAs, we undertook a biochemical and structural investigation of a putative xrRNA from TABV. We found that this xrRNA is indeed exoribonuclease resistant, and through solving the three-dimensional structure of the RNA to high resolution by x-ray crystallography, that it utilizes the characteristic ring-like fold. Some of the interactions found in the class 1 xrRNAs are also present in TABV, but other interactions are distinct when compared to previously characterized xrRNAs. These discoveries suggest a potential new subclass of xrRNAs within the *Flaviviridae*, which awaits confirmation once more members have been identified. The existence of a divergent flavivirus xrRNA structure related to previously-solved structures, but formed with different tertiary contacts, demonstrates how evolution can craft diverse RNA sequences to achieve a similar structural goal.

## RESULTS

### The TABV 3′ UTR contains an xrRNA dependent on long-range base pairs

Two regions in the 3′ UTR of TABV were identified as putative xrRNA1 and xrRNA2 (Ochsenreiter et al. 2019), however, they do not match the consensus sequence and secondary structure model of MBFV xrRNAs (**Sup. Fig. S1A**). The correct secondary structure of these two putative xrRNAs was ambiguous in that at least two plausible models could be proposed (**Sup. Fig. S1B**). One model more closely resembles the class 1 but appears to lack the sequences that form the tertiary interactions needed to form the ring-like structure. The other model is more similar to class 2, but neither model fully matches either class. This raises questions as to whether these divergent sequences in TABV are true xrRNAs, if they form the characteristic ring-like fold, and if so, how. Due to the 82% shared sequence identity between xrRNA1 and xrRNA2, we chose to focus on xrRNA1 for structural studies and construct design (**Sup. Fig. S1C)**.

We first determined whether the putative TABV xrRNA1 was exoribonuclease resistant using an established *in vitro* Xrn1 resistance assay. Briefly, we *in vitro* transcribed and purified RNA containing the wild-type (WT) sequence of the putative xrRNA1 from TABV with a 33-nucleotide endogenous “leader” sequence upstream of the putative xrRNA to allow Xrn1 to load and begin degradation. The RNA was challenged with recombinant Xrn1 and the reaction was resolved by denaturing gel electrophoresis. The results show processing of the full-length input RNA to a shorter resistant product, indicating an authentic xrRNA (**Fig. 2A, B**). We then mapped the 3′ stop site of Xrn1 using reverse transcription (RT). Previously characterized MBFV xrRNAs contained Xrn1 stop sites around 5 nucleotides upstream of the start of the P1 stem (Chapman et al. 2014b; Kieft et al. 2015; Akiyama et al. 2016; MacFadden et al. 2018) (**Sup. Fig. S2**). On TABV xrRNA1 the halt point was upstream of the resistant sequence, but because the structure was not yet solved, the relationship between the halt point and the structure was not known (**Sup. Fig. S1B)**.

**Figure 2.**
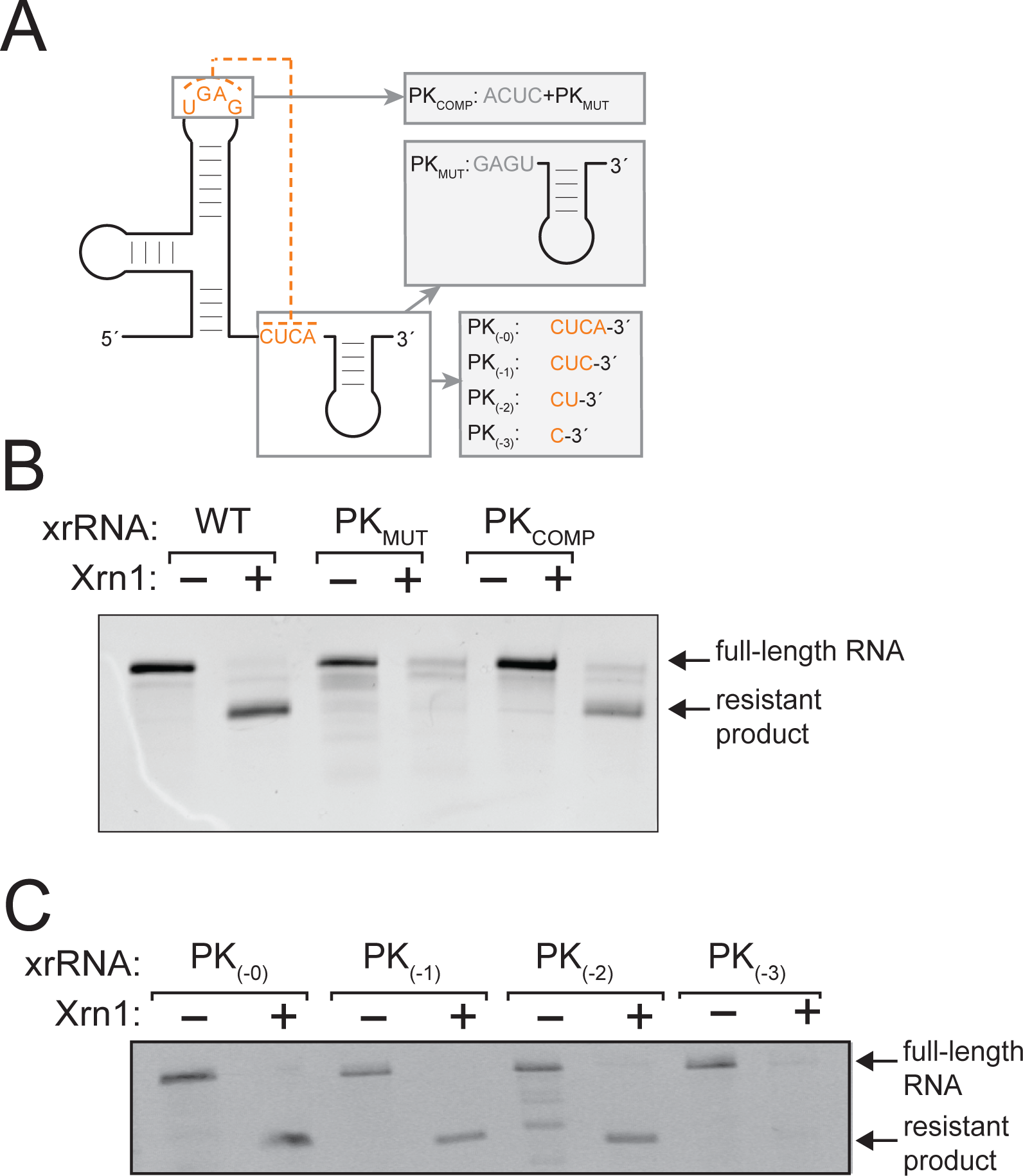
Functional need for a long-range pseudoknot in the TABV xrRNA. (A) Diagram of substitution mutations (PK_MUT_, PK_COMP_) and truncation mutations (PK_(−0)_, PK_(−1)_, PK_(−2)_, PK_(−3)_) used to test which elements are required for Xrn1 resistance. Full secondary structures of these mutants are contained in **Sup. Fig. 3A&B**. (B) Denaturing gel with results of Xrn1-resistance assay testing WT TABV RNA and two substitution mutants. Bands corresponding to full-length input RNA and the product of Xrn1 resistance are indicated. (C) Results of Xrn1 resistance assay with four truncation mutants.

Because all previously characterized xrRNA functionally require an important pseudoknot (PK) interaction (Funk et al. 2010; Silva et al. 2010; Chapman et al. 2014b; Kieft et al. 2015; Akiyama et al. 2016; MacFadden et al. 2018; Steckelberg et al. 2018a), we next generated and tested two mutant RNAs in addition to the wild-type (WT) to assess if this is also true in TABV xrRNA1. These were PK_MUT_, in which three bases in the 3′ end predicted to form long-range pairs were changed to their Watson-Crick reverse-complements, and PK_COMP_, which contained the same mutations as PK_MUT_ as well as compensatory mutations in the apical loop predicted to form the pseudoknot (**Fig. 2A, Sup. Fig. S3A**). PK_MUT_ was fully degraded, suggesting that mutation of these bases eliminates Xrn1 resistance, while PK_COMP_ showed restored ability to resist Xrn1. Together, these results establish the presence of an xrRNA in the 3′ UTR of TABV and show that long-range base pairs between the apical loop and downstream sequence are important for function.

To test the number of base pairs in the PK required for Xrn1 resistance and the need for sequence downstream of the PK, we next created a series of TABV xrRNA1 mutants by step-wise truncation of the 3′ end (PK_-0_, PK_-1_, PK_-2_, PK_-3_) (**Fig. 2A, Sup. Fig. S3B)**. Surprisingly, truncation of the 3′ end to a point where only two of the putative base-pairs can form still resulted in an xrRNA with some resistance to Xrn1 (**Fig. 2C)**. Further truncation to a single base-pair eliminated resistance. These results suggest that while the pseudoknot interaction is important for exoribonuclease resistance, other interactions within the TABV xrRNA stabilize the functional structure. Hence, like previously studied xrRNAs, the TABV sequence likely has a complex fold dependent on a variety of specific intramolecular contacts.

### The crystal structure of the TABV xrRNA reveals features differing from the MBFV xrRNA

To understand the structural basis of Xrn1 resistance by the TABV xrRNA, we solved the structure of the TABV xrRNA1 to 2.1 Å resolution by x-ray crystallography (**Fig. 3A, Sup. Table 1**). The crystallized TABV xrRNA1 is 52-nucleotides long, contains a GAAA tetraloop in P2, ends just after the PK, and retains Xrn1 resistance (**Fig. 3B**). The TABV xrRNA1 adopts a similar global conformation to previously solved flavivirus xrRNAs, with the distinctive ring-like fold encircling the 5′ end (**Fig. 3C**). In addition, the structure contains other features that are found in the MBFV xrRNA, including a base triple between C3, G25 and C39, Watson-Crick pairing between the 5′ end (S1) and the P1-P2-P3 junction nucleotides, and the aforementioned PK (**Fig. 3B**)(Kieft et al. 2015). Thus, overall the structure conforms to the global patterns expected of a flavivirus xrRNA and many of the tertiary interactions are maintained.

**Figure 3.**
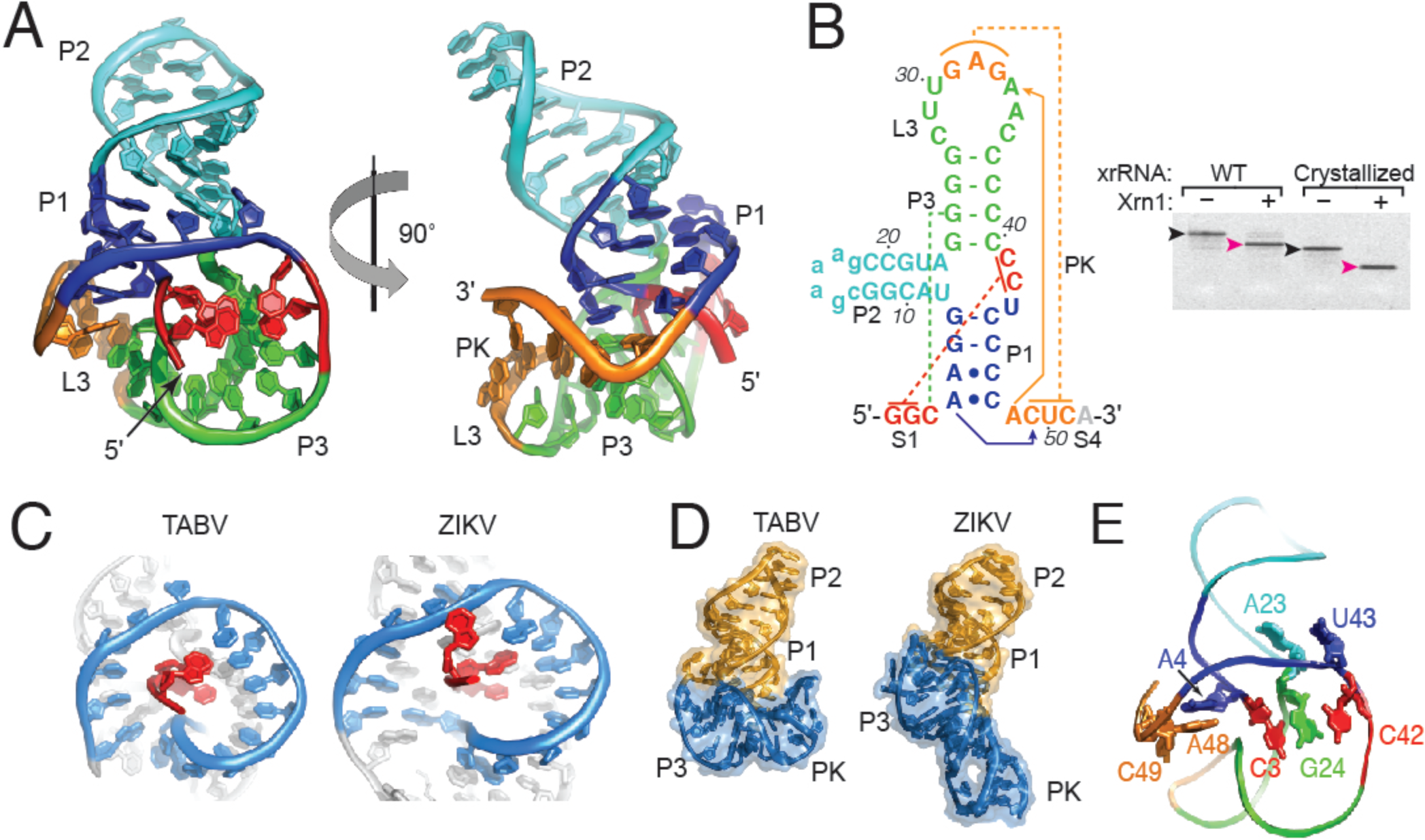
Crystal structure of TABV xrRNA. (A) X-ray crystal structure of the TABV xrRNA, with domains labeled and colored to match the secondary structure in panel (B). (B) Left: Secondary structure of the crystallized RNA. Lowercase: sequence altered from the native sequence. Dotted lines: tertiary interactions (red: internal pseudoknot; green: base triple; orange: long-range pseudoknot). Solid lines with arrows indicate two long-interactions involving As. Right: Xrn1 resistance assay of the native (WT) RNA and the crystallized RNA. Black arrow indicates input RNA, magenta arrow indicates protected fragment. (C) Ring-like structures of TABV and ZIKA compared side-by-side. The 5’ end is red, the ring is blue. (D) Full view of the TABV and ZIKV xrRNA structures side by side. The two distinct stacks are colored orange and blue in TABV, the corresponding regions are colored to match in ZIKV. (E) Cartoon of the backbone of the TABV structure highlighting the locations of sharp kinks in the backbone that give rise to distinct helical stacks: between C42 and U43, A23 and G24, C3 and A4, and A48 and C49.

Despite this similarly, the way in which the TABV xrRNA1 forms the structure is markedly different from other solved structures. Interestingly, the correct secondary structure of the xrRNA does not match any previously predicted possibilities (**Sup. Fig. S1B**). The three-way junction contains more nucleotides than was predicted, and the P1 stem consists of only two Watson-Crick base pairs with two adjacent A-C interactions as compared to five Watson-Crick pairs in the MBFV (**Fig. 3B, Sup. Fig. S1A**). Likewise, P3 has only four base-pairs, compared to 4-8 pairs in the MBFV consensus model. The stretch of pyrimidines on the 3’ side of P1 and P3 is shifted in register from what was predicted, and both the P1 and P3 stems are shorter than those in MBFV xrRNAs. Interestingly, in the TABV xrRNA, the U-A-U base triple found in all MBFV xrRNAs is replaced by an isosteric C^+^-G-C triple. This substitution was predicted based on mutational data using the xrRNA from the Cell fusing agent virus (MacFadden et al. 2018), the TABV xrRNA1 structure confirms the existence of this interaction. In addition, the putative xrRNA2 in the TABV 3’UTR can conform to this secondary structure (**Sup. Fig. S1C**). In addition, the mapped Xrn1 halt site is 5 nucleotides upstream of the base-pairs formed between the 5′ end (S1) and the P1-P2-P3 junction (**Sup. Fig. S2**), matching previously-characterized flavivirus xrRNAs (Chapman et al. 2014a; Akiyama et al. 2016).

To accommodate the conserved tertiary interactions within an RNA with shorter helices, the overall conformation of the TABV xrRNA has a distinct bend not observed in the structure of a MBFV xrRNA with longer helices (**Fig. 3D**). This bend in the TABV xrRNA structure gives rise to two separate helical stacks; one comprises P1 and P2, the other P3 and the L3 -S4 pseudoknot. In contrast, the previously-solved MBFV Zika virus xrRNA1 structure contains largely continuous base stacking throughout without a distinct sharp bend (Akiyama et al. 2016). The bent architecture of the TABV xrRNA1 is enabled by four distinct ‘breaks’ in base stacking where the backbone kinks (**Fig. 3E**). These lie between nucleotides A4-C3 (between S1 and P1), A23-G24 (P2 and P3), C42-U43 (in J123), and A48-C49 (immediately upstream of the L3-S4 pseudoknot) (**Sup. Fig. S4**). This bent-over conformation gives rise to a fold that, despite containing the same ring-like feature, is more compact than that of previously solved flavivirus xrRNAs. In addition, the crystallized TABV xrRNA1 lacks the P4 stem, which was present in the ZIKV xrRNA1 structure where it stacks directly on the PK. In TABV there is a stretch of U bases that are predicted to be unpaired between the PK and P4, hence the two are unlikely to stack in the same way. However, for xrRNAs tested thus far, the P4 element is not needed for Xrn1 resistance (Chapman et al. 2014b; Akiyama et al. 2016) (**Fig. 3B**).

### A complex set of interactions close the loop in the TABV xrRNA

The ring-like fold that encircles the 5′ end is important for exoribonuclease resistance, but in the TABV xrRNA1 the interactions that close the ring deviate dramatically from those in the MBFV. In Zika, for example, a single base extrudes from L3 to reverse Watson-Crick pair with a base in S4 that is sandwiched by base stacking on either side (**Fig. 4A**). Alignment of many MBFV class 1 xrRNAs and the structure of the xrRNA from Murray Valley encephalitis virus (MVEV) suggest this is a conserved feature (Chapman et al. 2014b; Akiyama et al. 2016). In contrast, in the TABV xrRNA1 the loop is closed by non-Watson-Crick interactions involving a network of hydrogen bonds mediated by metal ions and water (**Fig. 4B**), accompanied by the aforementioned kinks in the backbone (**Fig. 3E, Sup. Fig. S4**). Specifically, In the TABV xrRNA1, the P1 stem contains only two Watson-Crick base pairs, with two A-C interactions at its base. In the first A-C interaction, the exocyclic amine of A5 forms a single hydrogen bond to the carbonyl of C46 (**Fig. 4C**). The resolution of the crystal structure allows visualization of additional interactions involving these bases, formed by waters and metal ions. Specifically, functional groups on both the A5 Hoogsteen face and the Watson-Crick face of C46 make hydrogen bonds to waters, which are coordinated to a Mg^2+^ ion. This ion is localized by inner-sphere interactions to two phosphate oxygens (C3 and A4) near the location of a break in the base stacking (**Sup. Fig. S4A, S5A**). This Mg^2+^ also coordinates a water that is positioned to form a hydrogen bond with C47, which in turn is positioned to form a single hydrogen bond between its carbonyl and the exocyclic amine of A4 in the second A-C interaction. Thus, while only one hydrogen bond is formed in each A-C interaction, the waters and a metal ion creates a specific network of interactions that knit this region together.

**Figure 4.**
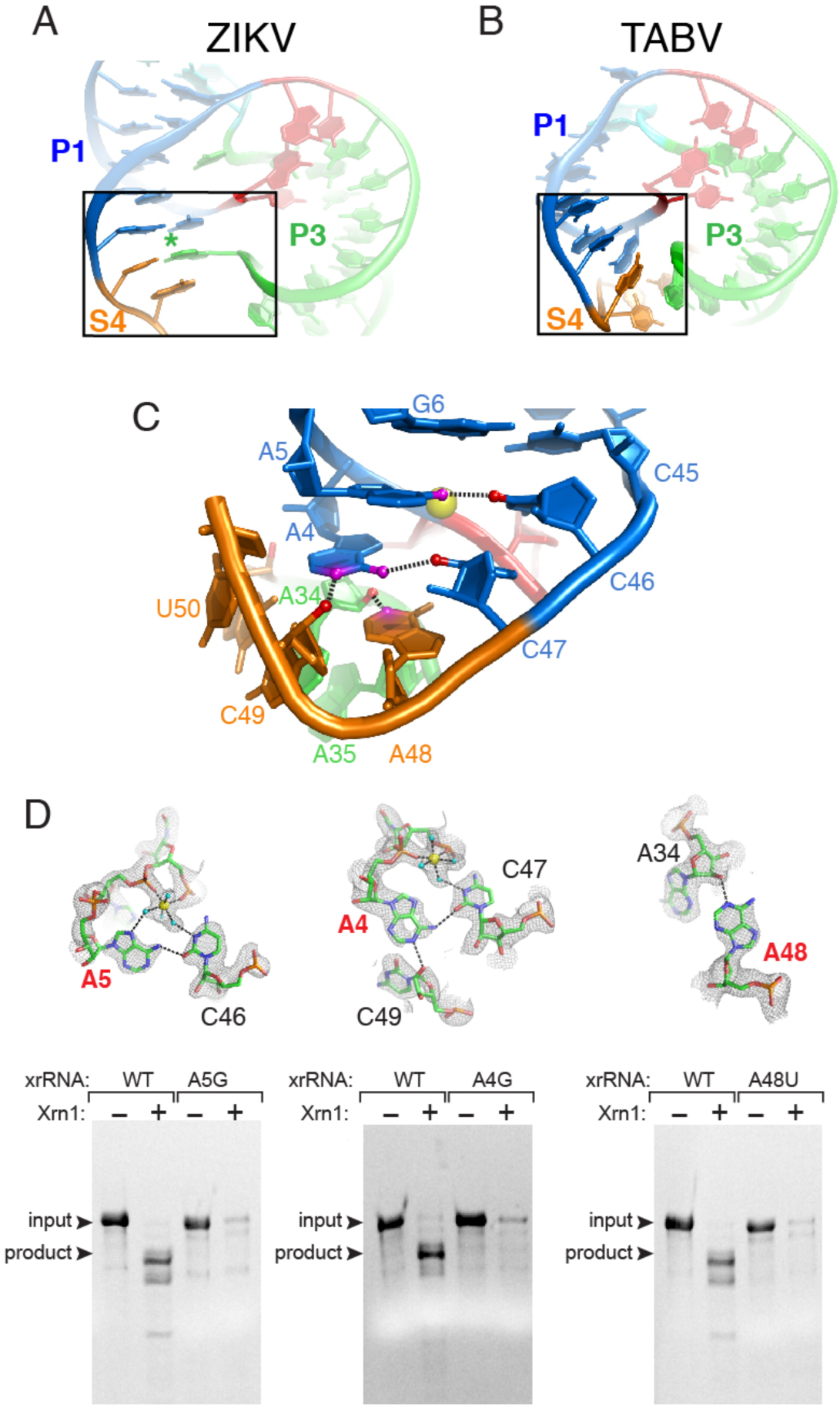
Interactions that close the ring in the TABV xrRNA. (A) Portion of the ZIKV xrRNA1 structure, highlighting contacts that close the ring-like element. (B) Portion of the TABV xrRNA1 crystal structure highlighting contacts that close the ring. (C) Close-up of panel (B) region, at a different orientation to show the network of interactions in the TABV xrRNA1. Colors of nucleotides match **Fig. 3A&B**. Nitrogen atoms and oxygen atoms involved in key hydrogen bonds are colored magenta and red, respectively. Dashed lines are hydrogen bonds. A Mg^2+^ ion is shown in yellow. (D) Top row: Close-up of three ring-closing contacts in TABV xrRNA1. Yellow spheres are Mg^2+^ ions, cyan balls are water molecules, and dotted lines represent hydrogen bonds or inner-sphere coordination of water to Mg^2+^. Electron density from a composite omit map is included. Nucleotides labeled in red were mutated and tested for Xrn1 resistance. Bottom row: Xrn1 resistance assays of WT and mutant xrRNAs. Bands corresponding to full-length input RNA and the product of Xrn1 resistance are indicated.

In addition to the A-C interactions, adenosines in this region form noncanonical interactions that appear structurally important. Specifically, the positioning of aforementioned A4 makes its Watson-Crick face available to form a hydrogen bond between its N1 and the 2’OH of C49. Thus, A4 interacts with both C47 and C49, forming a single hydrogen bond with each (**Fig. 4C**). The intervening nucleotide, A48 is adjacent to the A4-C47 pair and also forms a hydrogen bond between its N3 and the 2’OH of A34. The A48 and A4 thus stack on each other, but point in opposite directions, forming a pseudo-symmetric motif (**Fig. 4C, Sup. Fig. S4E**). These interactions close the ring; however, none match interactions observed in previous xrRNA structures and they create a more tightly packed structure than is observed in the xrRNA from ZIKV (**Fig. 4A**).

The TABV xrRNA structure suggests that the tertiary interactions described above are essential for Xrn1 resistance. If so, substitution of other nucleotides into some positions would break these interactions and negatively affect resistance. In particular, the adenosine residues in the P1 stem and S4 region appear to be structurally critical, therefore we generated mutant RNAs in which A4, A5, and A48 were replaced with guanosines (**Fig. 4D**). The A4G mutation was expected to eliminate or alter the A4-C46 interaction and also disrupt interactions with the Mg^2+^-coordinated waters. Likewise, A5G could have prevented the observed A5-C47 interaction from occurring, while also disrupting the formation of its A-minor interaction. Finally, A48G could have prevented the interaction that appears to be a key part of closing the ring. When subjected to Xrn1 degradation assays, all three A to G point mutants failed to exhibit resistance (**Fig. 4D**), demonstrating these bases have important roles in stabilizing the correct structure.

### Critical metal ions for structure and crystal packing

In addition to the Mg^2+^ ion described above, several other bound metal ions are positioned to play roles in stabilizing the exoribonuclease-resistant TABV xrRNA structure. One Mg^2+^ is found in the major groove of the interface between P3 and L3, where it is inner-sphere coordinated with a phosphate oxygen of A34 (**Sup. Fig. S5B**). Five waters are coordinated to this Mg^2+^, which in turn participate in a network of hydrogen bonds with adjacent nucleotides through base functional groups, phosphates and a 2’OH. One of these bonds is to the 2’OH of C3, which is in the third base in the C^+^-G-C formed between C3, G25, and C39. This Mg^2+^ ion-mediated network of interactions is associated with a narrowing of the major groove and may help stabilize the C^+^-G-C triple. Another Mg^2+^ ion is inner-sphere coordinated to a phosphate oxygen of C44, where it coordinates five water molecules. One of these is positioned to hydrogen bond to a phosphate oxygen of G2 at the location of a severe kink in the backbone and a break in base stacking (**Sup. Fig. S4D)**. In this location, the Mg^2+^ appears to help hold these regions in proximity while perhaps stabilizing the kink in the backbone. Several other Mg^2+^ ions and associated waters are also visible in the electron density, although their role in stabilizing the tertiary fold is not obvious. These may contribute to the overall compactness of the structure and likely reflect the high concentration of Mg^2+^ needed for crystallization. In addition, a single Na^+^ ion coordinated to six water molecules was also localized between backbone phosphates of G1 (near the 5’ end) and C36 (in P3); this ion was assigned based on the ion-water distances (Leonarski et al. 2017; Leonarski et al. 2019).

Two Mg^2+^ ions do not appear to be directly involved in structure stabilization but are interesting from a technical standpoint. The first is located at the interface between two RNA molecules in the crystal lattice, making inner-sphere coordination interactions with a phosphate oxygen of C47 on one molecule and to the phosphate oxygen of C13 on an adjacent molecule (**Sup. Fig. S5C)**. The other is found at a crystallographic special position (2-fold rotation axis) where it does not make any inner-sphere contact, but coordinated waters are able to hydrogen bond to both molecules (**Sup. Fig. S5D**). These ions likely stabilize the crystal.

### Diverged TABV xrRNA fails to tolerate critical MBFV xrRNA features

All MBFV xrRNAs contain a conserved cytosine in the 3-way junction that, when mutated to a G, eliminates the ability of the xrRNA block Xrn1, and this has become a mutation used in several contexts to disrupt xrRNA function and sfRNA formation (Chapman et al. 2014a; Chapman et al. 2014b; Kieft et al. 2015; Akiyama et al. 2016). Surprisingly, this C is not present in the TABV xrRNA1. An explanation for this is that the compact nature of the TABV ring creates a structure in which the C no longer fits. Specifically, the diameter of the MVEV and ZIKV xrRNAs rings are ∼30 Å and ∼31 Å respectively, while the TABV xrRNA is only 28 Å across (**Fig. 5A**). This difference is due to the aforementioned difference in the way the rings are closed, with the two MBFV using an extruded base to “reach out” and make a base-pair (**Fig. 4A, 5A**). This smaller ring also moves the 5’ end of the TABV xrRNA closer to P1 (**Fig. 5B**), such that it is involved in making ring-closing interactions through Mg^2+^ ion-mediated hydrogen bonds (**Fig. 4C, Sup. Fig. S5A**). The 5’ end does not do this in the MBFV xrRNAs. These seemingly subtle differences lead to the hypothesis that the two MBFV xrRNA structures have a pocket into which a C fits and makes contacts to part of the ring backbone, while the dimensions of the TABV xrRNA ring predict that this pocket is too small to accommodate a base without steric clash (**Fig. 5B**).

**Figure 5.**
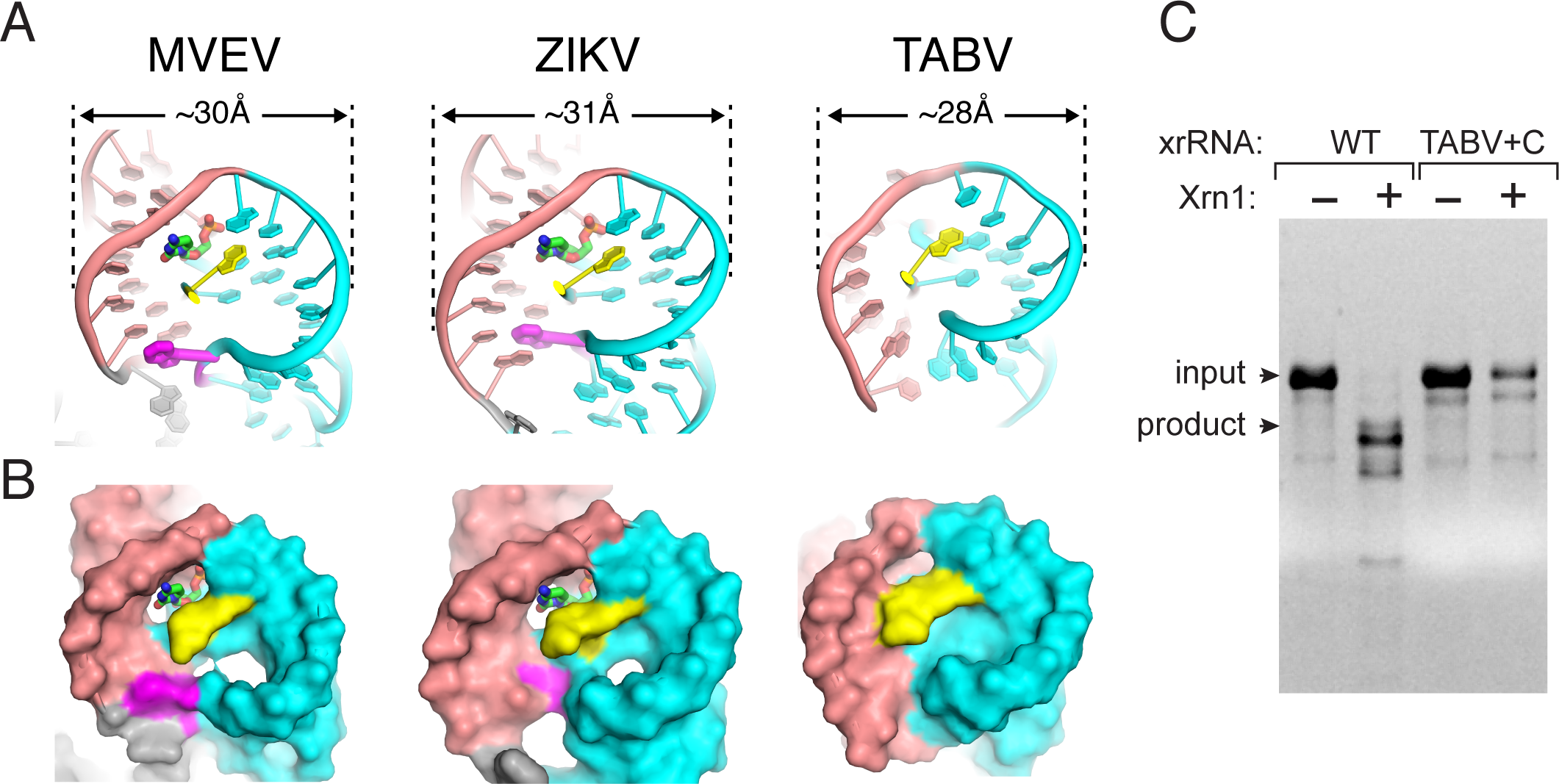
Differences in ring width mandate other changes in the structure. (A) Cartoon depiction of MVEV, ZIKV and TABV xrRNAs comparing the ring widths in Angstroms at their widest points. Salmon and teal define the two halves of the ring, yellow designates the 5′ end. Magenta indicates the nucleotide in MVEV and ZIKV that extrudes to reach across and make ring-closing contacts; there is no such nucleotide in TABV. A cytosine that is present in all MBFV is shown in MVEV and ZIKV, this is missing in TABV. (B) Same as panel (A), showing the solvent-accessible surface. The cytosine in MVEV and ZIKA fits in a pocket, this pocket is too small in TABV. (C) Denaturing gel with results of Xrn1 resistance assay performed on WT TABV xrRNA and TABV+C xrRNA, the latter has a cytosine inserted into the three-way junction where it is found in MBFV. Bands corresponding to full-length input RNA and the product of Xrn1 resistance are indicated.

To test the hypothesis that the TABV xrRNA1 structure could not accommodate a C in the position where it is always found in the MBFV xrRNAs, we created the TABV+C mutant by inserting a C after nucleotide 23 (**Sup. Fig. S3**), then analyzed this RNA for Xrn1 resistance. This mutation eliminates the ability of the TABV xrRNA1 to block degradation (**Fig. 5C**). Thus, this sequence element essential for resistance in MBFV is not tolerated in TBAV, suggestive of an evolutionary divergence in which certain interactions that stabilize the structure have been lost but been replaced by other interactions throughout the structure, revealing a different way to form a ring-like fold.

## DISCUSSION

The class 1 xrRNAs in the 3’UTRs of MBFV are defined by a distinct conserved pattern of co-varying bases, tertiary interactions, and primary sequence (MacFadden et al. 2018, Ochsenreiter, 2019 #169). xrRNAs of the divergent TABV lack these features, but also did not match the class 2 xrRNAs (Ochsenreiter et al. 2019). The high-resolution crystal structure of a functional xrRNA1 from TABV, presented here, shows that this RNA forms the characteristic ring-like conformation, but does so using a different set of tertiary interactions compared to other flavivirus xrRNAs (Chapman et al. 2014a; Akiyama et al. 2016; Steckelberg et al. 2018a; Steckelberg and Vicens et al. 2020). These findings have implications for the commonalities and differences in xrRNAs, for thinking of the features that drive function, and for understanding the distribution and evolution of xrRNAs in the *Flaviviridae* and beyond.

The fact that the TABV xrRNA forms the ring-like topology adds additional evidence that this fold is an important, elegant and perhaps widespread RNA structure-based strategy for blocking progression of 5’ to 3’ exoribonucleases. To date, this feature is found in all xrRNAs for which a three-dimensional structure has been solved (Chapman et al. 2014a; Akiyama et al. 2016; Steckelberg et al. 2018a; Steckelberg and Vicens et al. 2020). This includes not only those from flaviviruses but also those from plant-infecting viruses. However, examination of these structures shows that ring-like folds can be created using diverse strategies in which the secondary structures, sequences, and tertiary interactions that stabilize the conformation are different. A shared feature of all xrRNA structures solved to date is a pseudoknot interaction between an apical loop and downstream sequence (Chapman et al. 2014a; Akiyama et al. 2016; Steckelberg et al. 2018a; Steckelberg and Vicens et al. 2020). While common to all thus far, it is also important to note the presence of an RNA pseudoknot in a given RNA structure is not sufficient to confer resistance as it must exist within an overall structure in which the 5’ end passes through a distinctive continuous ring-like feature ∼15 nts long (Chapman et al. 2014a). Achieving this likely requires a specific folding pathway that is programmed into each xrRNA and that varies depending on the type (Kieft et al. 2015).

The TABV xrRNA structure reveals an unexpected number of tertiary interactions and features that deviate from the canonical Watson-Crick pairing. These include sharp backbone kinks and complex hydrogen bonding networks involving inner-sphere coordinated Mg^2+^ ions and water. The compact structure we observed is dependent on these interactions to a degree that is not obvious when just the secondary structure is viewed. This raises an important point: care must be taken when assigning ‘xrRNA-like’ status to elements that have secondary structures that superficially resemble authentic xrRNAs but do not form the requisite tertiary contacts. A three-way helical junction and likely pseudoknot interaction, all due to Watson-Crick pairing, are not sufficient to confer exoribonuclease resistance, illustrated here by the fact that the addition of a single C in a key location is sufficient to disrupt function **(Fig. 5C)**. Likewise, predictions of nuclease resistance based on predictions of thermodynamic stability of a predicted secondary structure are likely to be incorrect. Rather, xrRNA function depends on interactions that are currently difficult or impossible to predict from sequence or secondary structure information alone. More structural information from diverse xrRNAs are needed to enable robust predictions.

The differences between the TABV xrRNA and the previously-solved structures from the mosquito-borne flaviviruses are profound enough that they appear to be evolutionary distinct lineages. Changes, such as alteration in the length of helical elements, have been compensated for by other changes throughout the molecule. This illustrates how all of the parts must evolve together in such a way that the important topological elements are maintained even as the strategy of forming them changes. This point is directly illustrated by the presence or absence of an “extra” C-base in the three-way junction: it is required in the MBFV context, not tolerated in the TABV context. Again, this demands caution when assigning xrRNA-like status to uncharacterized RNAs based on the presence of certain hallmarks or superficial 2-D similarities. A difference in a single base in one part of the structure may require changes in other locations that are not readily predictable.

The TABV xrRNA, which has evolutionarily diverged from the mosquito-borne flavivirus xrRNAs, yet maintains function, now informs and motivates studies to look throughout the entire *Flaviviridae* family to see if other examples of TABV-like xrRNAs exist and how widespread they may be (Szucs et al. 2020). The success of such searches is greatly enhanced by the availability of correct secondary and tertiary structure information. Indeed, it seems likely that the secondary structure of some previously-suspected xrRNAs will need to be re-drawn and some lineages thought to not harbor xrRNAs may in fact have them. Finally, the increase in available structural information on diverse xrRNAs will inform efforts to find more examples throughout biology.

## METHODS

### Plasmid construction and cloning

To generate plasmids encoding the RNA sequences of interest, DNA templates were ordered as gene fragments (gBlocks) synthesized by Integrated DNA technologies (IDT) and cloned into pUC19 vector (NEB). Cloned plasmids were amplified in *E. coli* (DH5α) and verified by sequencing. Constructs for Xrn1 degradation assays contained the xrRNA sequence as well as 33 nucleotides of endogenous ‘leader sequence’ (UUAGCUUUGUUAGGUAGGGAAUUUUAAGAGAAA) to allow loading of the enzyme Xrn1. Below are the sequences of the TABV 3′ UTR as used in Xrn1 degradation assays with the T7 promoter underlined, the leader sequence in italics and minimal xrRNA highlighted in bold. Transcription with T7 RNA polymerase was started with 3 guanosine nucleotides (shown in lower case, not part of the endogenous viral sequence) for enhanced transcription efficiency. Superscript numbers within the TABV sequence denote the last nucleotide of TABV constructs PK_(−3)_, PK_(−2)_, PK_(−1)_, PK_(−0)_ and WT (in Fig. 1A and 1B), respectively. Bold nucleotides denote mutations.

TABV:

<u>TAATACGACTCACTATA</u>gggTTAGCTTTGTTAGGTAGGGAATTTTAAGAGAAAGGCAAGGTACGGATTA GCCGTAGGGGCTTGAGAACCCCCCCTCCCCAC^PK-3^T^PK-2^C^PK-1^A^PK-0^TTTTATTTCCTCTATGAGGAAG^WT^GTTAGGTTTGGGCAAGGTGCAG GTTAGCTGCAGGGGCTTGAAAAACCCCCCCCCCCATTCAAGACTTTTAGTGCATTAGTTTTAGGGTAAAGGGCAAGGTAGAGTGTTTCGCTCTAAGGGCGTAGGAACC

TABV PK MUT:

<u>TAATACGACTCACTATA</u>gggTTAGCTTTGTTAGGTAGGGAATTTTAAGAGAAAGGCAAGGTACGGATTA GCCGTAGGGGCTTGAGAACCCCCCCTCCCCA**GAGT**TTTTATTTCCTCTATGAGGAAG

TABV PK COMP:

<u>TAATACGACTCACTATA</u>gggTTAGCTTTGTTAGGTAGGGAATTTTAAGAGAAAGGCAAGGTACGGATTA GCCGTAGGGGCTT**CTC**AACCCCCCCTCCCCA**GAGT**TTTTATTTCCTCTATGAGGAAG

### RNA Preparation

DNA templates for *in vitro* transcription were amplified by PCR using custom DNA primers (IDT) and Phusion Hot Start polymerase (New England BioLabs). 2.5 mL transcription reactions were assembled using a 1000 μL PCR reaction, containing approximately 0.2 μM template DNA. This template DNA was combined with 6 mM each NTP, 60 mM MgCl_2_, 30 mM Tris pH 8.0, 10 mM DT, 0.1% spermidine, 0.1% Triton X-100, T7 RNA polymerase and 1 μL RNasin RNase inhibitor (Promega) and incubated overnight at 37°C. Next, inorganic pyrophosphates were removed through centrifugation. The reaction was ethanol precipitated and purified on a 7 M urea 8% denaturing polyacrylamide gel. Correct-sized RNA was cut from the gel and eluted in ∼50 mL of diethylpyrocarbonate (DEPC)-treated milli-Q filtered water (Millipore), washed, and concentrated using Amicon Ultra spin concentrators (Millipore).

### Protein expression and purification

The expression vector for *Kleuveromyces lactis* Xrn1 (Chang et al. 2011) (residues 1-1245) was a gift of Prof. Liang Tong at Columbia University and the expression vector for *Bdellovibrio bacteriovorus* RppH (ORF176/NudH/YgdP) (Messing et al. 2009) as a gift of Joel Belasco at NYU. Both recombinant proteins were 6XHis-tagged, expressed in *E. coli* BL21 cells and purified using Ni-NTA resin (Thermo), then followed by size exclusion with either a Superdex 75 or Superdex 200 column in an AKTA pure FPLC (GE Healthcare). The final protein product was stored in buffer containing 20 mM Tris pH 7.3, 300 mM NaCl, 1 mM DTT or 2 mM BME, and 10% glycerol (1 mMEDTA was added to the RNase J1 sample) at -80°C. The purity of the recombinant proteins was verified by SDS-PAGE and Coomassie staining

### 5’-3’ exoribonuclease degradation assay

For the degradation reaction, 5 μg RNA was resuspended in 50 μL 100 mM NaCl, 10 mM MgCl_2_, 50 mM Tris pH 7.5, 1 mM DTT. This reaction was incubated at 90°C for 3 min then 20°C for 5 min. 3 μL (0.5 μg/μL stock) RppH was added to the reaction to convert tri-phosphates on the 5’ end of RNA stands to mono-phosphates to allow Xrn1 loading, and then the reaction was split in two 25 μL reactions (+/- Xrn1 exoribonuclease). 1 μL of Xrn1 (0.8 μg/μL stock) was added to the Xrn1 positive half of the reaction, while a control sample of 1 μL of DEPC water was added to the Xrn1 negative reaction. All reactions were incubated at 37°C for 2 hours in a thermocycler. These degradation reactions were removed and resolved on a 7 M urea 8% denaturing polyacrylamide gel and stained with ethidium bromide.

### Reverse transcription mapping of exoribonuclease 1 (Xrn1) halt site

To determine the Xrn1 stop site at single-nucleotide resolution, 50 μg in vitro-transcribed RNA was processed using recombinant RppH and Xrn1 as described above (the reaction volume was scaled up to 400 μL, and 20 μL of each enzyme was used). The degradation reaction was resolved on a 7 M urea 8% polyacrylamide gel, then the Xrn1-resistant degradation product was cut from the gel and eluted passively overnight into RNase-free water. RNA was then washed and concentrated using Amicon Ultra spin concentrators (Millipore). After recovery, RNA was reverse-transcribed using Superscript III reverse transcriptase (Thermo) and a 6-FAM (6-fluorescein amidite)-labeled sequence-specific reverse primer (IDT) with an (A)20 -stretch at the 5′ end to allow cDNA purification with oligo(dT) beads. 10 μL RT reactions contained 1.2 pM RNA, 0.25 μL of 0.25 μM FAM-labeled reverse primer, 1 μL 5x First-Strand buffer, 0.25 μL of 0.1 M DTT, 0.4 μL of 10 mM dNTP mix, 0.1 μL Superscript III reverse transcriptase (200 U/μL) and were incubated for 1 hour at 50°C. To hydrolyze the RNA template after reverse transcription, 5 μL of 0.4 M NaOH was added and the reaction mix incubated at 90°C for 3 min, followed by cooling on ice for 3 min. The reaction was neutralized by adding 5 μL of acid quench mix (1.4 MNaCl, 0.57 M HCl, 1.3 M sodium acetate pH 5.2), then 1.5 μL oligo(dT) beads (Poly(A)Purist MAG Kit (Thermo)) were added and the cDNA purified on a magnetic stand according to the manufacturer’s instructions. The cDNA was eluted in 11 μL ROX-HiDi and analyzed on a 3500 Genetic Analyzer (Applied Biosystems) for capillary electrophoresis. A Sanger sequencing (ddNTP) ladder of the undigested RNA was analyzed alongside the degradation product as reference for band annotation.

### RNA crystallization and diffraction data collection

TABV xrRNA crystallization constructs varied in the length of P2 stem and the absence or presence of the P4 stem and loop. All constructs replaced the P2 loop with a ‘GAAA’ tetraloop. Six different constructs of TABV were screened for crystal formation, only one yielded crystals with diffraction suitable for structure determination. The sequence used for *in vitro* RNA transcription of this certain RNA was: 5′ - GCGAGTGAATTCtaatacgactcactatag**GGCAAGGTACGG**CGAAAG**CCGTAGGGGCTTGAGAACCCCCCCTCCCCACTCA***gggcggcatggtcccagcctcctcgctggcgccgcctgggcaacatgcttcggcatggcgaatgggacc*GGATCC GCGAT – 3′. The T7 polymerase promoter and one extra G added for transcription efficiency is shown in lowercase, the xrRNA sequence is shown in bold, with the portio that was changed for crystallization underlined. The hepatitis delta ribozyme used to generate a homogenous 3′ end is shown in italics.

Template DNA to generate RNA for crystallization was amplified by PCR directly from a gBlock template. Transcription reactions were conducted as outlined in “*In vitro* transcription”, then ribozyme cleavage was enhanced at the end of transcription by adding MgCl_2_ (final conc. 120 mM) and incubating for 5 minutes at 65°C. RNA was then purified on a 7 M urea 8% polyacrylamide gel as describe in the “*In vitro* transcription” section. RNA was refolded in reactions containing 5 μg/μL RNA, 30 mM HEPES (pH 7.5), 20 mM MgCl_2_ and 100 mM KCl at 65°C for 2 minutes and then centrifuged at full speed for 2 minutes. Crystal Screens I and II and Natrix I and II (all from Hampton Research) were used for the initial RNA screen at room temperature in sitting-drop vapor diffusion crystal trays (Axygen). An initial hit in 0.2 M Mg^2+^ chloride hexahydrate, 0.1 M HEPES sodium (pH 7.5), and 30% v/v Polyethylene glycol 400 provided three crystals which were flash-frozen in liquid nitrogen for x-ray diffraction.

Diffraction data were collected at Advanced Light Source Beamline 4.2.2 using ‘shutterless’ collection at the Iridium L-III edge (1.0972 Å) at 100°K. 180° datasets were collected with 0.2° oscillation images. Data were indexed, integrated, and scaled using XDS. The buffer containing the initial hit never yielded any other crystals, so one non-derivatized crystal was used to solve the structure by molecular replacement. To accomplish this, we first identified a search model through a search of the Protein Data Bank (PDB) for RNA structures with sequence similar to any in the TABV xrRNA. We identified an RNA hairpin with a sequence identical to the GAAA loop in P2 of the TABV xrRNA, specifically nucleotides 8-17 (chain P), residues 8-17 (chain Q), residues 26-35 (chain P) and residues 57-66 (chain P) from the L1 Ribozyme Ligase circular adduct structure (PDB ID: 2OIU) (Robertson and Scott 2007). We aligned the four fragments taken from two copies in the asymmetric unit to generate an ensemble, which was used in Phenix to determine an initial molecular replacement solution.

### Model building and refinement

The initial map from molecular replacement was very incomplete. Iterative rounds of model building and refinement (simulated annealing, Rigid-body, B-factor refinement, phase combination) using COOT (Emsley et al. 2010) and Phenix (v.1.16_3549) were used to generate a complete model of the RNA on MAC OS (Liebschner et al. 2019). Magnesium and sodium ions were assigned based on geometry (Leonarski et al. 2017; Leonarski et al. 2019). Waters were placed during late stages of refinement using autopicking in Phenix. Crystal diffraction data, phasing, and refinement statistics are contained in **Sup. Table S1**.

## DATA DEPOSITION

The coordinates have been deposited in the Protein Data Bank with the accession code XXXX (to be supplied prior to publication)

## ACKNOWLEDGEMENTS

The authors thank current and former Kieft Lab members for thoughtful discussions and technical assistance. We also thank Drs. Mark Sterken and Gorban Pijlman for alerting us to putative xrRNAs in TABV. This work was supported by NIH grants R35GM118070 and R01AI133348 (JSK), and T32AI052066 (MJS). RAJ is supported by an NIH diversity supplement to grant R35GM118070. A-LS was funded by a Deutsche Forschungsgemeinschaft Fellowship STE2509/201. Beamline 4.2.2 at the Advanced Light Source is partially funded by the NIH (P30GM124169-01) and operated under contract with the U.S. DOE(DE-AC02-05CH11231). The University of Colorado Anschutz Medical Campus x-ray crystallography facility is supported by the NIH (P30CA046934 and S10OD012033).

## SUPPLEMENTARY MATERIALS

**Supplementary Table 1.**
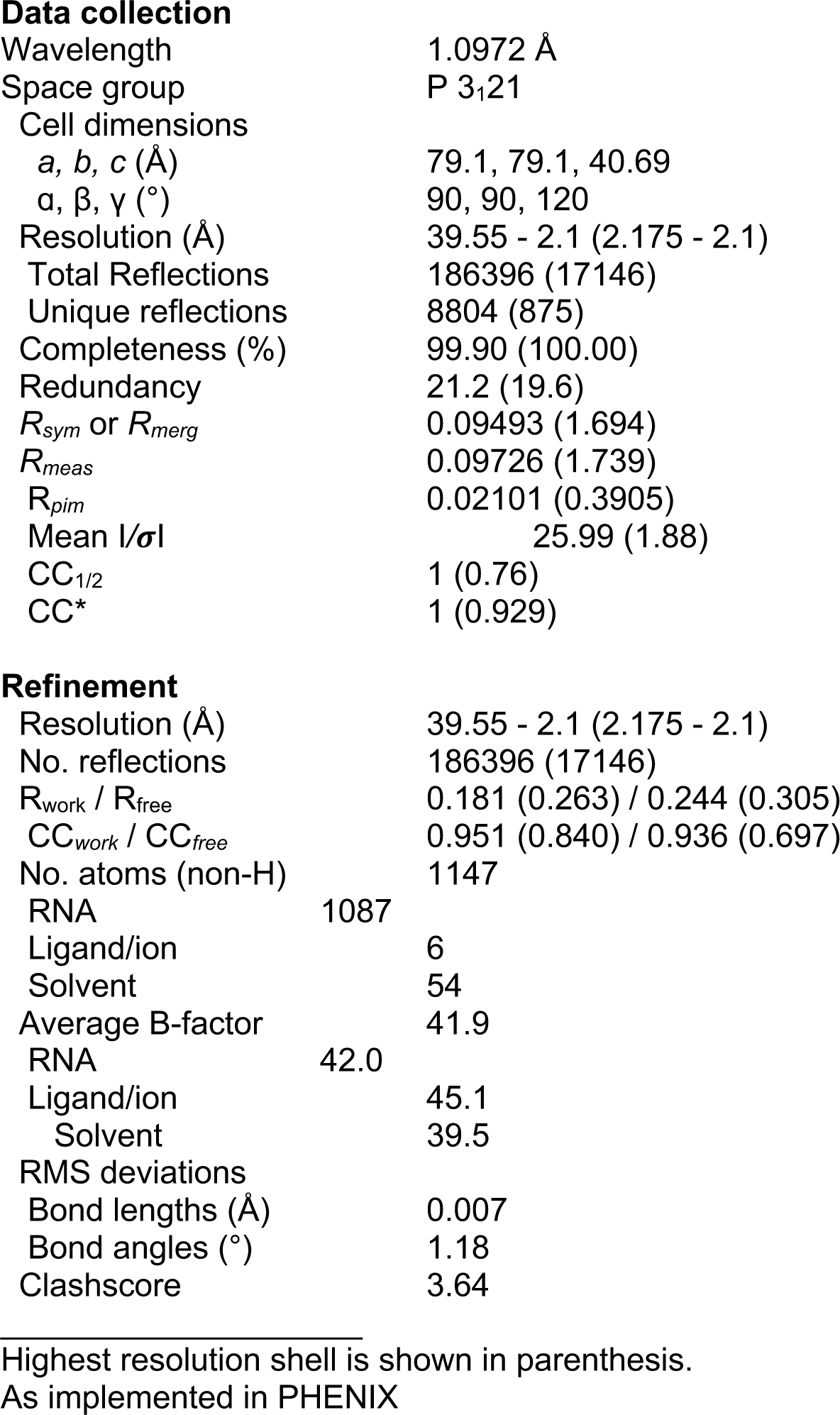
Crystallographic data collection and refinement statistics. Highest resolution shell is shown in parenthesis. As implemented in PHENIX

|  |  |
| --- | --- |
| Wavelength | 1.0972 Å |
| Space group | P 3<sub>1</sub>21 |
| Cell dimensions |  |
| *a*, *b*, *c* (Å) | 79.1, 79.1, 40.69 |
| $\alpha$, $\beta$, $\gamma$ (°) | 90, 90, 120 |
| Resolution (Å) | 39.55 - 2.1 (2.175 - 2.1) |
| Total Reflections | 186396 (17146) |
| Unique reflections | 8804 (875) |
| Completeness (%) | 99.90 (100.00) |
| Redundancy | 21.2 (19.6) |
| $R_{sym}$ or $R_{merg}$ | 0.09493 (1.694) |
| $R_{meas}$ | 0.09726 (1.739) |
| $R_{pim}$ | 0.02101 (0.3905) |
| Mean $I/\sigma$ | 25.99 (1.88) |
| CC<sub>1/2</sub> | 1 (0.76) |
| CC* | 1 (0.929) |
| **Refinement** |  |
| Resolution (Å) | 39.55 - 2.1 (2.175 - 2.1) |
| No. reflections | 186396 (17146) |
| $R_{work}$ / $R_{free}$ | 0.181 (0.263) / 0.244 (0.305) |
| CC<sub>work</sub> / CC<sub>free</sub> | 0.951 (0.840) / 0.936 (0.697) |
| No. atoms (non-H) | 1147 |
| RNA | 1087 |
| Ligand/ion | 6 |
| Solvent | 54 |
| Average B-factor | 41.9 |
| RNA | 42.0 |
| Ligand/ion | 45.1 |
| Solvent | 39.5 |
| RMS deviations |  |
| Bond lengths (Å) | 0.007 |
| Bond angles (°) | 1.18 |
| Clashscore | 3.64 |

**Supplementary Figure S1.**
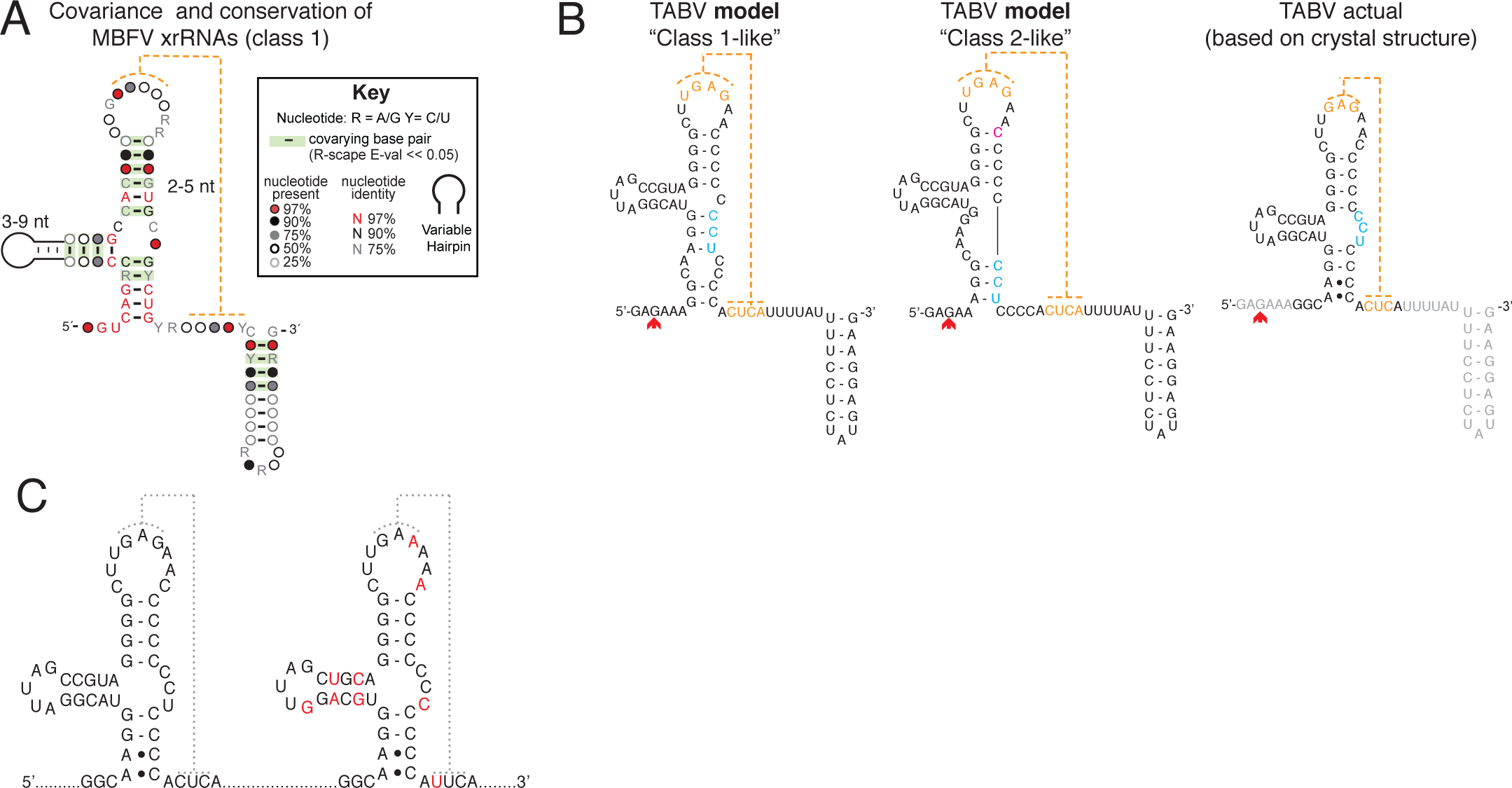
Putative TABV xrRNA secondary models. (A) Covariance diagram of the class 1xrRNAs found in mosquito-borne flaviviruses (Szucs et al. 2020). This diagram shows the pattern of base-pairs and conservation that defines this class of xrRNA. The legend for this diagram is inset. (B) Two possible secondary structures of the TABV xrRNA. The first resembles class 1, while the second is more like class 2. However, neither matches the consensus diagram of panel (A) and conserved sequences known to be critical for folding the class 1 are not present. At right is the experimentally determined secondary structure of the TABV xrRNA1 based on the crystal structure (grey sequences flanking the secondary structure were not present in the crystallized RNA). Red arrows denote the Xrn1 stop site. Nucleotides in blue are provided to emphasize differences between the structural diagrams. Pseudoknot interaction is in orange. (C) Secondary structure models of both TABV xrRNA1 and xrRNA2 in the 3′ UTR. The xrRNA2 is predicted to match the 3D-based 2D structure of xrRNA1. xrRNA1 was tested and is the source of the RNA crystallized here. The downstream putative xrRNA has not been tested for resistance, it contains nine nucleotide differences compared to the upstream (red).

**Supplementary Figure S2.**
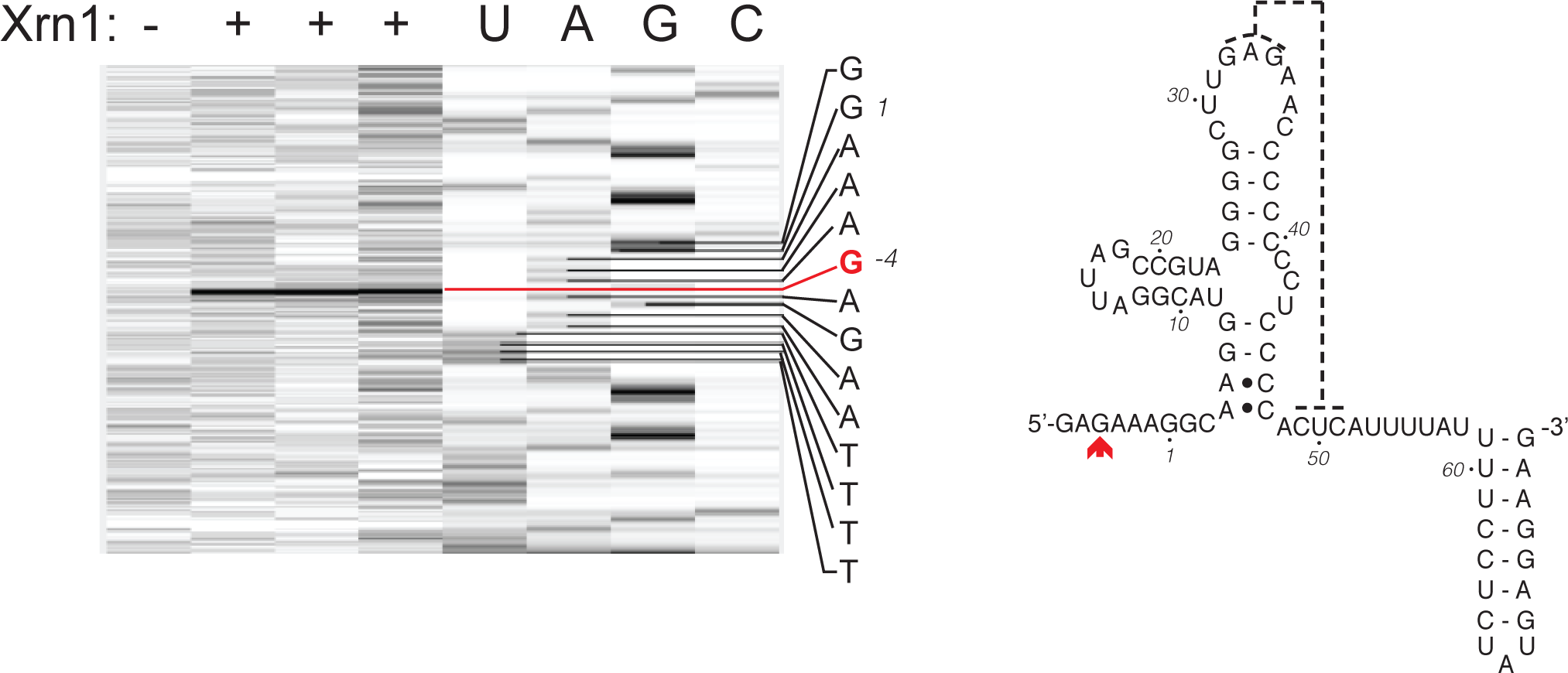
Mapping the Xrn1 halt site on Stop-site analysis of the TABV xrRNA. (A) To initially characterize the TABV xrRNA structure, we mapped the 3′ stop site of Xrn1 using reverse transcription (RT). Briefly, undigested input TABV xrRNA and three samples of TABV digested by Xrn1 were reverse transcribed using a fluorescently labeled primer, then analyzed by capillary electrophoresis and compared against dideoxy-containing sequencing reactions. Full length TABV xrRNA appears in the first column, while the next three display the digested TABV sequence after incubation with Xrn1 nuclease. The thick band shows the 5’ end of the RNA remaining after Xrn1 treatment, and thus the halt point of the enzyme. The sequence of the xrRNA is shown to the right with the red G and line indicating the halt point of the enzyme. (B) The secondary structure of the TABV xrRNA1, derived from the crystal structure, is shown. The halt point of Xrn1 is indicated with a red arrow. Numbering matches the crystallized RNA.

**Supplementary Figure S3.**
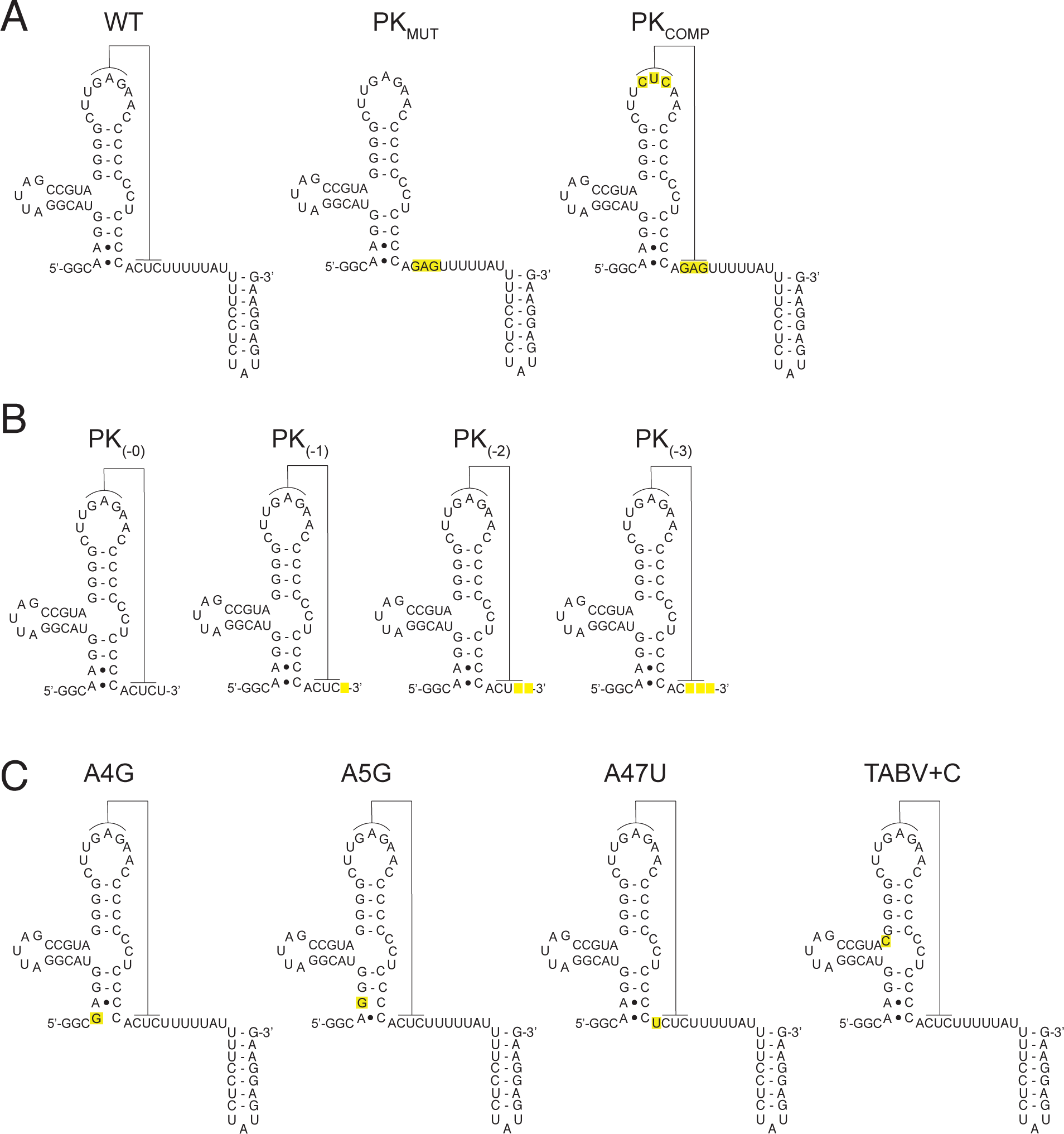
Mutants used in this study. (A) Secondary structure diagrams of the three RNAs used to test the presence of a pseudoknot interaction, results are in **Figure 2B**. Yellow indicates locations of substitution mutations. (B) Secondary structure of truncation mutants used to identify the minimum length of TABV xrRNA, results are in **Figure 2C**. Yellow indicates the missing nucleotides. (C) Secondary structure of mutant RNAs used to determine the importance of A bases at critical points in the structure. Yellow indicates the location of substitution mutations. Results are in **Figure 4D**. (D) Mutant with a C base inserted into the three-way junction as it is found in MBFV. Yellow indicates the location for the insertion. Results are in **Figure 5C**.

**Supplementary Figure S4.**
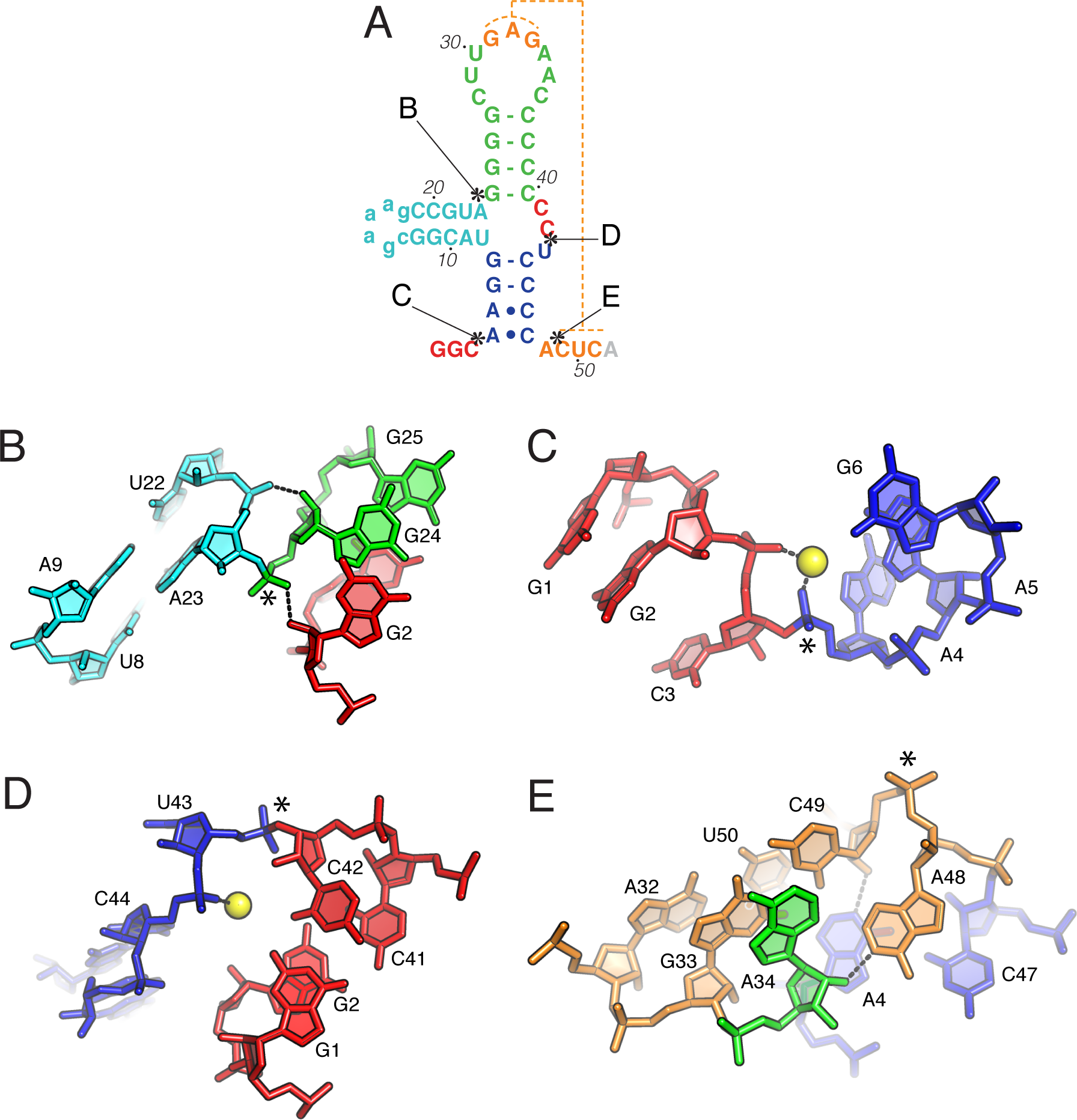
Four distinct breaks in base stacking at kinked positions in the backbone. (A) Secondary structure of the TABV xrRNA with asterisks denoting the location of the four kinks in the structure, which are labeled as they appear in this figure. (B) Detailed view of the kink between A23 and G24. (C) Detailed view of the kink between C3 and A4, in which inner sphere coordination to a Mg^2+^ (yellow sphere) by two adjacent phosphates plays a key role. (D) Detailed view of the kink between U43 and C42, which is also facilitated by Mg^2+^ (yellow sphere). (E) Detailed view of the kink between A48 and C49. In this area, note the hydrogen bonds (dashed lines) between N1 of A48 and the 2’OH of A34, and between the N1 of A4 and the 2′OH of C49. In all panels, dashed lines indicate important hydrogen bonds.

**Supplementary Figure S5.**
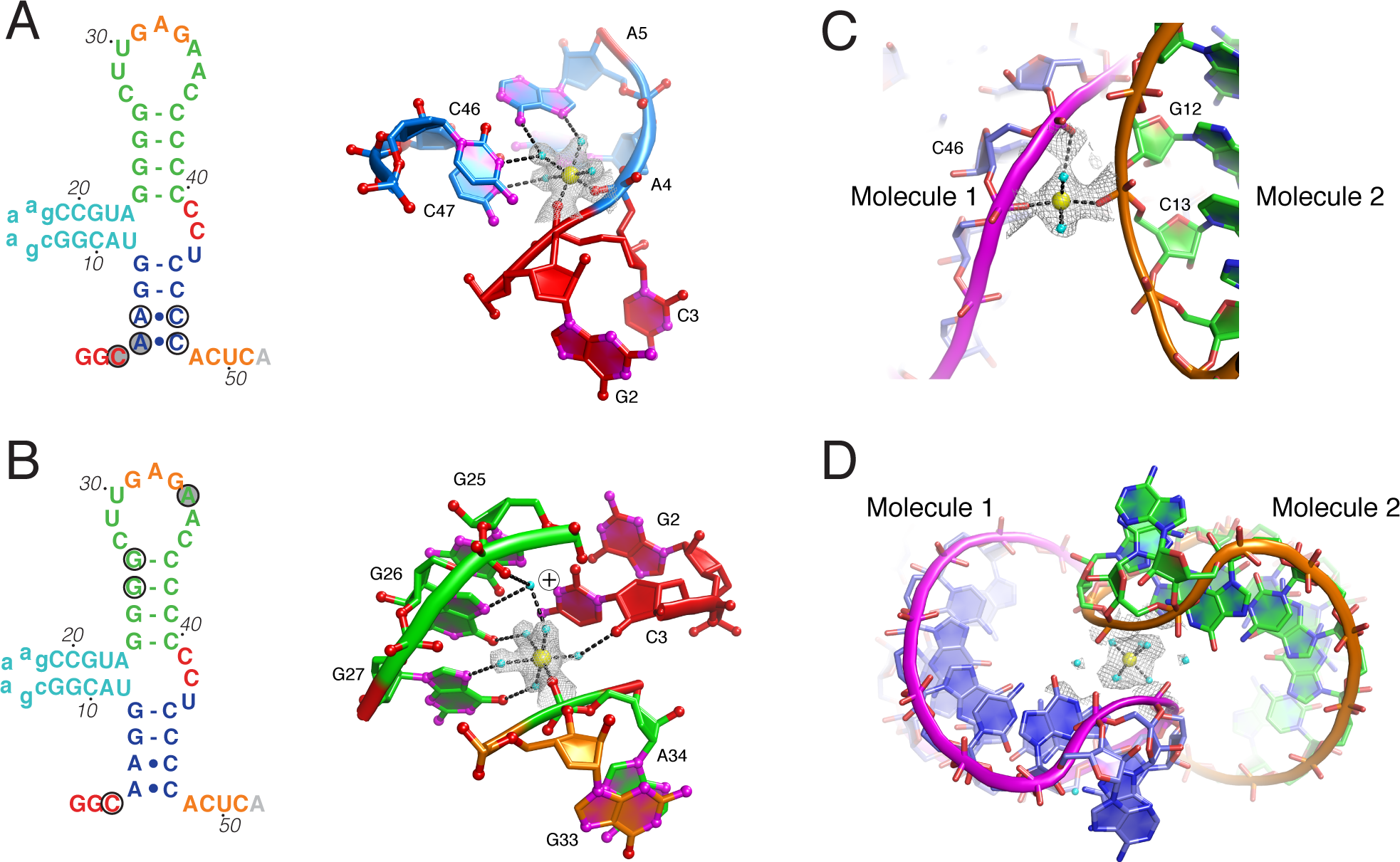
Mg^<u>2</u>+^ ions contribute to the folding of the TABV xrRNA1 and crystal packing. (A) Left: Secondary structure of the TABV xrRNA with circled nucleotides denoting those that interact with a localized Mg^2+^ ion. Shaded nucleotides are making inner-sphere coordination, while open circles designate nucleotides that hydrogen bond to with waters that are coordinated to the Mg^2+^ ion. Right: The Mg^2+^ ion (yellow sphere) is shown surrounded by density from a composite omit map. Dashed lines indicate hydrogen bonds or coordination to the Mg^2+^. In the RNA, the bases are colored to match the secondary structure, and oxygen atoms are colored red, nitrogen atoms are colored magenta. (B) Left: Secondary structure indicating the location of another bound Mg^2+^, using the same scheme as panel (A). Right: Bound Mg^2+^ is shown as in panel (A). The circled plus sign indicates the location of a protonation on the C base, suggested by its participation in the C^+^-G-C base triple. (C) Close-up view of a unique Mg^2+^ ion that mediates a crystal contact between two molecules in the crystal lattice. The ion makes inner-sphere coordination to phosphates in two different RNA molecules in the crystal. (D) A Mg^2+^ ion bound at a crystallographic special position (on a twofold rotation axis).

## REFERENCES

Akiyama BM, Laurence HM, Massey AR, Costantino DA, Xie X, Yang Y, Shi PY, Nix JC, Beckham JD, Kieft JS. 2016. Zika virus produces noncoding RNAs using a multi-pseudoknot structure that confounds a cellular exonuclease. Science 354: 1148–1152.

Bidet K, Dadlani D, Garcia-Blanco MA. 2014. G3BP1, G3BP2 and CAPRIN1 are required for translation of interferon stimulated mRNAs and are targeted by a dengue virus non-coding RNA. PLoS Pathog 10: e1004242.

Brinton MA, Basu M. 2015. Functions of the 3’ and 5’ genome RNA regions of members of the genus Flavivirus. Virus Res 206: 108–119.

Chang JH, Xiang S, Xiang K, Manley JL, Tong L. 2011. Structural and biochemical studies of the 5’-->3’ exoribonuclease Xrn1. Nat Struct Mol Biol 18: 270–276.

Chang RY, Hsu TW, Chen YL, Liu SF, Tsai YJ, Lin YT, Chen YS, Fan YH. 2013. Japanese encephalitis virus non-coding RNA inhibits activation of interferon by blocking nuclear translocation of interferon regulatory factor 3. Vet Microbiol 166: 11–21.

Chapman EG, Costantino DA, Rabe JL, Moon SL, Wilusz J, Nix JC, Kieft JS. 2014a. The structural basis of pathogenic subgenomic flavivirus RNA (sfRNA) production. Science 344: 307–310.

Chapman EG, Moon SL, Wilusz J, Kieft JS. 2014b. RNA structures that resist degradation by Xrn1 produce a pathogenic Dengue virus RNA. Elife 3: e01892.

Clarke BD, Roby JA, Slonchak A, Khromykh AA. 2015. Functional non-coding RNAs derived from the flavivirus 3’ untranslated region. Virus Res 206: 53–61.

de Borba L, Villordo SM, Marsico FL, Carballeda JM, Filomatori CV, Gebhard LG, Pallares HM, Lequime S, Lambrechts L, Sanchez Vargas I et al. 2019. RNA Structure Duplication in the Dengue Virus 3’ UTR: Redundancy or Host Specificity? mBio 10.

de Lamballerie X, Crochu S, Billoir F, Neyts J, de Micco P, Holmes EC, Gould EA. 2002. Genome sequence analysis of Tamana bat virus and its relationship with the genus Flavivirus. J Gen Virol 83: 2443–2454.

Donald CL, Brennan B, Cumberworth SL, Rezelj VV, Clark JJ, Cordeiro MT, Freitas de Oliveira Franca R, Pena LJ, Wilkie GS, Da Silva Filipe A et al. 2016. Full Genome Sequence and sfRNA Interferon Antagonist Activity of Zika Virus from Recife, Brazil. PLoS Negl Trop Dis 10: e0005048.

Emsley P, Lohkamp B, Scott WG, Cowtan K. 2010. Features and development of Coot. Acta Crystallogr D Biol Crystallogr 66: 486–501.

Fauquet CM, Mayo MA, Maniloff J, Desselberger U, Ball LA. 2005. Virus Taxonomy: VIIIth Report of the International Committee on Taxonomy of Viruses. Elsevier Science.

Fields BN, Knipe DM, Howley PM. 2013. Fields virology. Wolters Kluwer/ Lippincott Williams & Wilkins Health, Philadelphia, PA, USA.

Filomatori CV, Carballeda JM, Villordo SM, Aguirre S, Pallares HM, Maestre AM, Sanchez-Vargas I, Blair CD, Fabri C, Morales MA et al. 2017. Dengue virus genomic variation associated with mosquito adaptation defines the pattern of viral non-coding RNAs and fitness in human cells. PLoS Pathog 13: e1006265.

Funk A, Truong K, Nagasaki T, Torres S, Floden N, Balmori Melian E, Edmonds J, Dong H, Shi PY, Khromykh AA. 2010. RNA structures required for production of subgenomic flavivirus RNA. J Virol 84: 11407–11417.

Goertz GP, van Bree JWM, Hiralal A, Fernhout BM, Steffens C, Boeren S, Visser TM, Vogels CBF, Abbo SR, Fros JJ et al. 2019. Subgenomic flavivirus RNA binds the mosquito DEAD/H-box helicase ME31B and determines Zika virus transmission by Aedes aegypti. Proc Natl Acad Sci U S A 116: 19136–19144.

Iwakawa HO, Mizumoto H, Nagano H, Imoto Y, Takigawa K, Sarawaneeyaruk S, Kaido M, Mise K, Okuno T. 2008. A viral noncoding RNA generated by cis-element-mediated protection against 5’->3’ RNA decay represses both cap-independent and cap-dependent translation. J Virol 82: 10162–10174.

Jones CI, Zabolotskaya MV, Newbury SF. 2012. The 5’ --> 3’ exoribonuclease XRN1/Pacman and its functions in cellular processes and development. Wiley Interdiscip Rev RNA 3: 455–468.

Junglen S, Korries M, Grasse W, Wieseler J, Kopp A, Hermanns K, Leon-Juarez M, Drosten C, Kummerer BM. 2017. Host Range Restriction of Insect-Specific Flaviviruses Occurs at Several Levels of the Viral Life Cycle. mSphere 2.

Kieft JS, Rabe JL, Chapman EG. 2015. New hypotheses derived from the structure of a flaviviral Xrn1-resistant RNA: Conservation, folding, and host adaptation. RNA Biol 12: 1169–1177.

Leonarski F, D’Ascenzo L, Auffinger P. 2017. Mg2+ ions: do they bind to nucleobase nitrogens? Nucleic Acids Res 45: 987–1004.

Leonarski F 2019. Nucleobase carbonyl groups are poor Mg(2+) inner-sphere binders but excellent monovalent ion binders-a critical PDB survey. RNA 25: 173–192.

Liebschner D, Afonine PV, Baker ML, Bunkoczi G, Chen VB, Croll TI, Hintze B, Hung LW, Jain S, McCoy AJ et al. 2019. Macromolecular structure determination using X-rays, neutrons and electrons: recent developments in Phenix. Acta Crystallogr D Struct Biol 75: 861–877.

MacFadden A, O’Donoghue Z, Silva P, Chapman EG, Olsthoorn RC, Sterken MG, Pijlman GP, Bredenbeek PJ, Kieft JS. 2018. Mechanism and structural diversity of exoribonuclease-resistant RNA structures in flaviviral RNAs. Nat Commun 9: 119.

Manokaran G, Finol E, Wang C, Gunaratne J, Bahl J, Ong EZ, Tan HC, Sessions OM, Ward AM, Gubler DJ et al. 2015. Dengue subgenomic RNA binds TRIM25 to inhibit interferon expression for epidemiological fitness. Science 350: 217–221.

Maruyama SR, Castro-Jorge LA, Ribeiro JM, Gardinassi LG, Garcia GR, Brandao LG, Rodrigues AR, Okada MI, Abrao EP, Ferreira BR et al. 2014. Characterisation of divergent flavivirus NS3 and NS5 protein sequences detected in Rhipicephalus microplus ticks from Brazil. Mem Inst Oswaldo Cruz 109: 38–50.

Messing SA, Gabelli SB, Liu Q, Celesnik H, Belasco JG, Pineiro SA, Amzel LM. 2009. Structure and biological function of the RNA pyrophosphohydrolase BdRppH from Bdellovibrio bacteriovorus. Structure 17: 472–481.

Michalski D, Ontiveros JG, Russo J, Charley PA, Anderson JR, Heck AM, Geiss BJ, Wilusz J. 2019. Zika virus noncoding sfRNAs sequester multiple host-derived RNA-binding proteins and modulate mRNA decay and splicing during infection. J Biol Chem 294: 16282–16296.

Moon SL, Anderson JR, Kumagai Y, Wilusz CJ, Akira S, Khromykh AA, Wilusz J. 2012. A noncoding RNA produced by arthropod-borne flaviviruses inhibits the cellular exoribonuclease XRN1 and alters host mRNA stability. RNA 18: 2029–2040.

Moon SL, Dodd BJ, Brackney DE, Wilusz CJ, Ebel GD, Wilusz J. 2015. Flavivirus sfRNA suppresses antiviral RNA interference in cultured cells and mosquitoes and directly interacts with the RNAi machinery. Virology 485: 322–329.

Naeve CW, Trent DW. 1978. Identification of Saint Louis encephalitis virus mRNA. J Virol 25: 535–545.

Nagarajan VK, Jones CI, Newbury SF, Green PJ. 2013. XRN 5’-->3’ exoribonucleases: structure, mechanisms and functions. Biochim Biophys Acta 1829: 590–603.

Ng WC, Soto-Acosta R, Bradrick SS, Garcia-Blanco MA, Ooi EE. 2017. The 5’ and 3’ Untranslated Regions of the Flaviviral Genome. Viruses 9.

Ochsenreiter R, Hofacker IL, Wolfinger MT. 2019. Functional RNA Structures in the 3’UTR of Tick-Borne, Insect-Specific and No-Known-Vector Flaviviruses. Viruses 11.

Pijlman GP, Funk A, Kondratieva N, Leung J, Torres S, van der Aa L, Liu WJ, Palmenberg AC, Shi PY, Hall RA et al. 2008. A highly structured, nuclease-resistant, noncoding RNA produced by flaviviruses is required for pathogenicity. Cell Host Microbe 4: 579–591.

Pompon J, Manuel M, Ng GK, Wong B, Shan C, Manokaran G, Soto-Acosta R, Bradrick SS, Ooi EE, Misse D et al. 2017. Dengue subgenomic flaviviral RNA disrupts immunity in mosquito salivary glands to increase virus transmission. PLoS Pathog 13: e1006535.

Robertson MP, Scott WG. 2007. The structural basis of ribozyme-catalyzed RNA assembly. Science 315: 1549–1553.

Schnettler E, Sterken MG, Leung JY, Metz SW, Geertsema C, Goldbach RW, Vlak JM, Kohl A, Khromykh AA, Pijlman GP. 2012. Noncoding flavivirus RNA displays RNA interference suppressor activity in insect and Mammalian cells. J Virol 86: 13486–13500.

Schnettler E, Tykalova H, Watson M, Sharma M, Sterken MG, Obbard DJ, Lewis SH, McFarlane M, Bell-Sakyi L, Barry G et al. 2014. Induction and suppression of tick cell antiviral RNAi responses by tick-borne flaviviruses. Nucleic Acids Res 42: 9436–9446.

Schuessler A, Funk A, Lazear HM, Cooper DA, Torres S, Daffis S, Jha BK, Kumagai Y, Takeuchi O, Hertzog P et al. 2012. West Nile virus noncoding subgenomic RNA contributes to viral evasion of the type I interferon-mediated antiviral response. J Virol 86: 5708–5718.

Selisko B, Wang C, Harris E, Canard B. 2014. Regulation of Flavivirus RNA synthesis and replication. Curr Opin Virol 9: 74–83.

Shi M, Lin XD, Chen X, Tian JH, Chen LJ, Li K, Wang W, Eden JS, Shen JJ, Liu L et al. 2018. The evolutionary history of vertebrate RNA viruses. Nature 556: 197–202.

Silva PA, Pereira CF, Dalebout TJ, Spaan WJ, Bredenbeek PJ. 2010. An RNA pseudoknot is required for production of yellow fever virus subgenomic RNA by the host nuclease XRN1. J Virol 84: 11395–11406.

Slonchak A, Hugo LE, Freney ME, Hall-Mendelin S, Amarilla AA, Torres FJ, Setoh YX, Peng NYG, Sng JDJ, Hall RA et al. 2020. Zika virus noncoding RNA suppresses apoptosis and is required for virus transmission by mosquitoes. Nat Commun 11: 2205.

Slonchak A, Khromykh AA. 2018. Subgenomic flaviviral RNAs: What do we know after the first decade of research. Antiviral Res 159: 13–25.

*Steckelberg A-L, *Vicens Q, Costantino DA, Nix JC, Kieft JS. 2020. The crystal structure of a Polerovirus exoribonuclease-resistant RNAshows how diverse sequences are integrated into a conserved fold. bioRxiv doi: 101101/20200430070631 *joint first authors

Steckelberg AL, Akiyama BM, Costantino DA, Sit TL, Nix JC, Kieft JS. 2018a. A folded viral noncoding RNA blocks host cell exoribonucleases through a conformationally dynamic RNA structure. Proc Natl Acad Sci U S A 115: 6404–6409.

*Steckelberg AL, *Vicens Q, Kieft JS. 2018b. Exoribonuclease-Resistant RNAs Exist within both Coding and Noncoding Subgenomic RNAs. mBio 9. *joint first authors

Szucs MJ, Nichols PJ, Jones RA, Vicens Q, Kieft JS. 2020. A new subclass of exoribonuclease resistant RNA found in multiple Flaviviridae genera. bioRxiv

Takeda H, Oya A, Hashimoto K, Yasuda T, Yamada MA. 1978. Association of virus specific replicative ribonucleic acid with nuclear membrane in chick embryo cells infected with japanese encephalitis virus. J Gen Virol 38: 281–291.

Walker SE, Zhou F, Mitchell SF, Larson VS, Valasek L, Hinnebusch AG, Lorsch JR. 2013. Yeast eIF4B binds to the head of the 40S ribosomal subunit and promotes mRNA recruitment through its N-terminal and internal repeat domains. RNA 19: 191–207.

Wengler G, Wengler G, Gross HJ. 1978. Studies on virus-specific nucleic acids synthesized in vertebrate and mosquito cells infected with flaviviruses. Virology 89: 423–437.

Yeh SC, Pompon J. 2018. Flaviviruses Produce a Subgenomic Flaviviral RNA That Enhances Mosquito Transmission. DNA Cell Biol 37: 154–159.

